# Impaired SOX17 Expression Causes Endothelial Dysfunction and Pulmonary Arterial Hypertension by Insufficient Suppression of RUNX1

**DOI:** 10.64898/2026.05.14.725187

**Authors:** Bedia Akosman, Moon Jung Choi, Yamini Sharma, Mandy Pereira, Young Eun Lee, Eui Young So, Allyson Sherman Roe, Navneet Singh, Anthony M. Reginato, Corey E. Ventetuolo, Martin R. Wilkins, Lan Zhao, Christopher J. Rhodes, James R. Klinger, Olin D. Liang

## Abstract

Genome-wide association studies have identified rare and common mutations associated with increased risk of pulmonary arterial hypertension (PAH), but the mechanism by which impaired SOX17 expression increases PAH risk is not known. Notably, SOX17 plays a critical role in endothelial identity during development by suppressing RUNX1 through binding to its promoter and directing stem and progenitor cells toward an endothelial rather than a hematopoietic cell fate. RUNX1 functions as a key regulator of myeloid differentiation, aberrant angiogenesis and adverse cardiac remodeling. Previously, we found that RUNX1 inhibition reverses pulmonary hypertension (PH) in multiple animal models. Here, we hypothesize that impaired expression of SOX17 in PAH leads to endothelial cell (EC) dysfunction by failing to suppress RUNX1.

**METHODS:** Human pulmonary artery endothelial cells (HPAECs) with stable SOX17 CRISPR/Cas9 knockout or RUNX1 overexpression were generated and examined for endothelial and hematopoietic gene expression, proliferation, migration, apoptosis, and angiogenesis. Immortalized lymphoblastoid cell lines (LCLs) from PAH patients with SOX17 mutations and healthy controls were reprogrammed into induced pluripotent stem cells (iPSCs) and differentiated into ECs. The effect of RUNX1 inhibition on Sugen/hypoxia-PH was examined in rats, SOX17 enhancer knockout (SOX17enhKO) mice, and Cdh5-CreERT2;Runx1(flox/flox);SOX17enhKO triple transgenic mice. SOX17 and RUNX1 expression were analyzed in peripheral blood samples from PAH patients (n=359).

**RESULTS:** HPAECs with SOX17 deletion or RUNX1 overexpression exhibited decreased expression of EC markers, enhanced proliferation and migration, defective angiogenesis, and decreased apoptosis. RUNX1 siRNA knockdown or RUNX1 inhibition by Ro5-3335 partially restored the endothelial properties in SOX17 KO HPAECs. ECs differentiated from SOX17 mutant PAH patient iPSCs exhibited upregulated RUNX1 expression and loss of endothelial identity, which was also partially restored by RUNX1 siRNA or Ro5-3335. In addition, SOX17enhKO mice had increased RUNX1 expression and susceptibility to Sugen/hypoxia-induced PH (SuHx-PH). Treatment with RUNX1 inhibitors or inducible endothelial-specific deletion of RUNX1 rescued SuHx-PH susceptibility in SOX17enhKO mice. RUNX1 inhibitors Ro5-3335 and Ro24-7429 also reversed SuHx-PH in wild-type rats. In addition, plasma RUNX1 expression was higher in PAH patients lacking detectable SOX17 expression than in patients with detectable SOX17 expression.

**CONCLUSIONS:** Impaired SOX17 expression increases the risk of PAH through insufficient suppression of RUNX1, leading to pulmonary endothelial dysfunction. RUNX1 inhibition mitigates PH associated with SOX17 deficiency and may represent a novel therapeutic strategy for PAH, especially those with rare or common SOX17 mutations.

## INTRODUCTION

Pulmonary arterial hypertension (PAH) is a type of PH characterized by a severe obliterative vasculopathy of the distal pulmonary circulation ^1–3^ that leads to marked increases in pulmonary vascular resistance resulting in severe limitations in functional capacity, right heart failure, and death. Median survival in untreated patients is only 3 years ^4^. Although rare, it occurs in 0.5-10% percent of patients with HIV infection, portal hypertensin, congenital heart disease, connective tissue diseases, and methamphetamine use ^4, 5^. In addition to PAH, PH occurs frequently in chronic heart and lung diseases such as diastolic heart failure, pulmonary fibrosis, and COPD, and is associated with similar increases in morbidity and mortality ^6, 7^. Despite the development of over a dozen drugs, representing 6 different classes, in the last 30 years, there is no effective therapy for this devastating disease. Currently available medications produce only moderate improvement in pulmonary arterial pressure (PAP) and 6-minute walking distance (6MWD) and none are curative. The 5-year survival rate even with modern therapy remains around 50% ^8, 9^. Thus, new therapies are needed that directly target pathogenic mechanisms responsible for the disease.

Genetic mutations are responsible for the majority of cases of familial PAH and an increasing number of cases of sporadic PAH. Several large genome-wide association studies (GWAS) have identified mutations in SOX17 with an increased risk of PAH. The SOX (SRY-related HMG-box) family of transcription factors that plays an important role in the development of the cardiovascular system 13 and angiogenesis 14, 15 due to its critical role in driving stem and progenitor cells to an endothelial rather than a hematopoietic cell fate 16 17. Rare missense and frameshift mutations in the SOX17 gene occur in about 1% of all PAH patients and 3% of PAH patients with congenital heart disease (CHD). These patients exhibit PAH at an earlier age and have more severe pulmonary vascular disease than patients with idiopathic PAH or heritable PAH caused by other mutations. SOX17 mutant patients exhibit severe obliteration of distal pulmonary vessels, loss of segmental distribution, corkscrewed vessels, and abnormal capillary blush ^10^. In addition to rare mutations in the gene, numerous single nucleotide variants (SNVs) in SOX17 have also been associated with PAH. These SNVs involve two independent signals in the enhancer region 100-200 kb upstream of the SOX17 gene. Risk alleles at both signals are common, such that 59% of PAH patients and 46% of controls, were homozygous for the risk allele at both signals. Endothelial specific knockout of SOX17 causes PH in adult mice ^11, 12^, and previous studies by our group demonstrate that SOX17enhancer knockout (SOX17enhKO), in which the signal 1 region containing common SOX17 SNVs associated with increased risk of PAH is deleted, develop more severe hypoxic PH and are more susceptible to Sugen/hypoxia PH (SuHx-PH) than wild-type (WT) mice ^13^. Together these findings demonstrate the important role that SOX17 plays in PAH pathogenesis. However, the mechanism by which impaired SOX17 expression increases the risk and severity of PAH is not understood.

During development, SOX17 plays a major role in driving stem and progenitor cells toward an endothelial fate by suppressing the expression of Runt-related transcription factor 1 (RUNX1). RUNX1 is a member of the core-binding factor (CBF) family of transcription factors that is indispensable for the establishment of definitive hematopoiesis in vertebrates via endothelial to hematopoietic transition (EHT) ^14–16^, hematopoietic transformation of adult bone marrow-derived progenitor cells, and myeloid differentiation ^17–19^. In earlier studies, we demonstrated that gene expression of RUNX1 is increased in circulating CD34+CD133+ progenitor cells in patients with PAH compared to healthy volunteers, and that a selective inhibitor of RUNX1 reverses established PH in two rodent models of PAH ^20^. We also showed that tissue specific deletion of *RUNX1* in adult endothelium or myeloid cells prevents SuHx-PH in mice ^21^.

SOX17 suppresses RUNX1 expression through specific binding sites upstream from the RUNX1 promotor ^22^. Small perturbations in SOX17 levels are accompanied by increased levels of RUNX1 hematopoietic fate ^23–25^. We hypothesized that the increased risk and severity of PH associated with impaired SOX17 expression is due to insufficient suppression of RUNX1. To test this hypothesis, we examined RUNX1 expression and endothelial function in human pulmonary artery endothelial cells (HPAECs) after SOX17 knockout and in iPSC-derived endothelial cells (iPSC-ECs) generated from PAH patients with SOX17 mutations. We also examined the expression of RUNX1 in the lung and bone marrow of SOX17enhKO mice during the development of PH. We\ then examined ability of RUNX1 inhibition restore endothelial function in vitro and prevent PH in SOX17enhKO mice. We found that reduced SOX17 expression was associated with increased RUNX1 levels. Both SOX17 deficiency and RUNX1 upregulation impaired endothelial identity and promoted endothelial dysfunction, and RUNX1 inhibition reversed these changes in vitro and reversed PH in vivo. Together, our findings suggest that SOX17/RUNX1 signaling plays an important role in the pathogenesis of PH and may be novel targets for the treatment of PAH.

## MATERIALS AND METHODS

### Generation of stable SOX17 knockout HPAECs and RUNX1 overexpressing HPAECs

Human pulmonary arterial endothelial cells (HPAECs; Lonza, Cat# CC-2530) were cultured in Endothelial Growth Medium-2 (EGM-2; Lonza, Cat# CC-3162) at 37°C in a humidified incubator with 5% CO₂. To generate stable SOX17 CRISPR/Cas9 knockout (KO) HPAECs, a single-guide RNA targeting the human SOX17 gene (5′-GTAGTACACGTGAAGGGCGC-3′) was cloned into the lentiCRISPRv2 vector (Addgene plasmid #52961). To generate stable RUNX1 overexpression HPAECs, full-length human RUNX1 cDNA (OriGene, Cat# SC123977) was cloned into the pHAGE-EF1α-IRES-blast vector from our lab. High titer lentiviruses were produced at Brown University Legorreta Cancer Center Cell & Genome Engineering Shared Resource. HPAECs were transduced at a multiplicity of infection (MOI) of 10 through spinfection, which was performed in the presence of 8 µg/ml polybrene (Sigma-Aldrich, Cat# TR-10003) at 900 g and 32°C for 2 h. The culture medium was replaced 24 h post-transduction, and puromycin antibiotic selection was initiated 5 days post-transduction with a concentration of 0.7 µg/ml. Cells between passages 3 and 6 were used for all experiments.

### RUNX1 siRNA downregulation and small molecule compound Ro5-3335 inhibition in vitro

RUNX1 knockdown was performed using siRNA targeting human RUNX1 (Dharmacon/Horizon Discovery, ON-TARGETplus SMARTpool, Cat# L-003926-00-0005). A non-targeting control siRNA pool served as the negative control. Electroporation was carried out using the Neon Transfection System (Thermo Fisher Scientific) with the R Buffer according to the manufacturer’s protocol. Briefly, 1 × 10e6 HPAECs were resuspended in the Neon R Buffer containing 50 nM siRNA, aspirated into the Neon 100 µl tip, and electroporated using the following parameters: 1150 V, 30 ms pulse width, 2 pulses. For pharmacological RUNX1 inhibition, the cells were treated with the small molecule inhibitor Ro5-3335 ^26^ (Sigma-Aldrich, Cat# 21-950-610MG) at final concentrations of 10 µM or 25 µM. Cells were treated for 24 h for RNA isolation or 48 h for protein isolation, with vehicle controls receiving an equivalent amount of DMSO.

### Reprogramming of lymphoblastoid cell lines to induced pluripotent stem cells

The National Biological Sample and Data Repository for PAH (NIH PAH Biobank) has biological samples, clinical data, and genetic data from approximately 2900 patients with WHO Group 1 PAH including 8 patients with missense or frameshift SOX17 mutations predicted to be deleterious (Supplementary Table S1). In collaboration with the Biobank PI Dr. William Nichols (Cincinnati Children’s Hospital Medical Center, Cincinnati, OH), we have obtained immortalized lymphoblastoid cell lines (LCLs) from these patients and age-and sex-matched control PAH patients without SOX17 mutations and reprogrammed them into iPSCs. Age- and sex-matched healthy control LCLs were purchased from Creative Biolabs (Shirley, NY).

LCLs were maintained in RPMI 1640 medium (Gibco, Cat# 11875-093) supplemented with 15% fetal bovine serum (FBS; Gibco, Cat# 26140079) until reprogramming. For each nucleofection reaction, 2 × 10e6 LCLs were resuspended in 100 µl of R buffer from the NEON Nucleofection Kit (Thermo Fisher Scientific, Cat# MPK10096) and nucleofected with 2.5 µg of reprogramming plasmids: pCE-hOCT3/4 (Addgene plasmid #41813), pCE-hSK (Addgene plasmid #41814), pCE-hUL (Addgene plasmid #41855), pCE-mp53DD (Addgene plasmid #41856), and pCXB-EBNA1 (Addgene plasmid #41857). Nucleofection was performed using the NEON Transfection System (Thermo Fisher Scientific, Cat# MPK5000) with the following parameters: 1300V, 10 ms pulse width, and a 3 pulses. Following nucleofection, cells were allowed to recover overnight in 2 ml of RPMI 1640 medium supplemented with 15% FBS. On the following day, half of the cell suspension was transferred to a hESC-qualified Matrigel-coated well (Corning, Cat# 354277) of a 6-well TC-treated plate containing 1 ml of complete ReproTeSR reprogramming medium (STEMCELL Technologies, Cat# 05926). Media changes were performed according to the following schedule: at days 3 and 5, 750 µl of ReproTeSR medium was added; at days 7 and 9, 1 ml of medium was replaced with fresh ReproTeSR; at day 11, a complete medium change was performed with ReproTeSR. At days 13 and 15, 1 ml of medium was replaced with mTeSR Plus medium (STEMCELL Technologies, Cat# 05825), followed by complete medium changes with mTeSR Plus every other day until iPSC colonies were ready for picking.

### Characterization of LCL-derived iPSCs

#### Western blot analysis of pluripotency markers

Pluripotency of iPSC clones was assessed by Western blot using the StemLight™ iPS Cell Reprogramming Antibody Kit (Cat# 9092, Cell Signaling Technology), including antibodies against OCT4, SOX2, NANOG, c-MYC, KLF4, and LIN28A. β-actin was used as a loading control.

#### Immunocytochemistry for pluripotency markers

Immunocytochemistry was performed using the PSC 4-Marker Immunocytochemistry Kit (Invitrogen, Cat# A24881) according to the manufacturer’s instructions. iPSC colonies (approximately 70-80% confluency) were fixed, permeabilized, and blocked prior to incubation with primary antibodies against SSEA, OCT4, SOX2, and TRA-1-60 at 4°C for 3 h. After washing, cells were incubated with secondary antibody for 1 h at room temperature and counterstained with NucBlue™ (DAPI). Stained colonies were imaged using an Olympus CKX53 phase contrast fluorescence microscope (Evident Scientific, Waltham, MA).

#### Trilineage differentiation

Trilineage differentiation was performed using the STEMdiff™ Trilineage Differentiation Kit (STEMCELL Technologies, Cat# 05230) according to the manufacturer’s instructions. For lineage validation, cells were fixed with 4% paraformaldehyde, permeabilized with 0.3% Triton X-100 in 1% BSA/PBS, and blocked in the same buffer. Endoderm and ectoderm markers were detected using anti-FOXA2/HNF3β (D56D6, Alexa Fluor 555, 1:100) and anti-NESTIN (10C2, Alexa Fluor 488, 1:100) antibodies (Cell Signaling Technology), respectively. Mesoderm differentiation was assessed using anti-Brachyury (R&D Systems, Cat# AF2085, 10 µg/ml), followed by incubation with Alexa Fluor 568-conjugated donkey anti-goat secondary antibody (1:500). Primary antibody incubation was performed overnight at 4°C. All samples were counterstained with DAPI and imaged using a fluorescence microscope.

#### Karyotype analysis

Karyotype analysis of iPSC lines was performed by STEMCELL Technologies using standard G-banding cytogenetic methods.

### Differentiation of endothelial cells from PAH patient LCL-derived iPSCs

Endothelial cells (ECs) were generated from PAH patient and healthy control LCL-derived iPSCs using two complementary approaches. In the first method, iPSCs were differentiated into ECs using the STEMdiff™ Endothelial Differentiation Kit (STEMCELL Technologies, Cat# 08005) according to the manufacturer’s instructions. Following differentiation, cells were stained with CD144 (VE-cadherin; Miltenyi Biotec, 1:50, Cat# 130-123-688) and CD31 (PECAM-1; Miltenyi Biotec, 1:50, Cat# 130-110-807) antibodies, and double-positive populations were isolated by fluorescence-activated cell sorting (FACS). Sorted cells were subsequently cultured using the STEMdiff™ Endothelial Expansion Kit (STEMCELL Technologies, Cat# 100-1218). In the second method, we utilized a modified version of a previously published chemical induction protocol ^27^, and iPSCs were maintained and differentiated upon reaching approximately 70-80% confluency. Differentiation was initiated by switching cells to RPMI 1640 medium containing B-27 Supplement Minus Insulin (Gibco, Cat# A1895601), and treating cells with CHIR99021 (STEMCELL Technologies, Cat# 72052) (6 µM for 48 h followed by 3 µM for another 48 h) to induce mesoderm formation. Cells were then exposed to vascular endothelial growth factor VEGF; 50 ng/ml, PeproTech, Cat# 100-20) and basic fibroblast growth factor ((bFGF; 10 ng/ml, PeproTech, Cat#100-18B) for 7 days in RPMI 1640 medium containing B-27 Supplement Minus Insulin. Cells were subsequently stained with anti-CD144 and anti-CD31 antibodies and subjected to FACS to isolate double-positive (CD144+/CD31+) ECs. Sorted cells were cultured in endothelial growth medium (EGM-2; Lonza, Cat# CC-3162) on gelatin-coated plates for expansion. The chemical induction method yielded a greater number of ECs with more consistent morphology and was therefore used for the majority of downstream experiments.

### RT² Profiler PCR Array analysis

Total RNA was isolated using the Trizol/Chloroform method. cDNA synthesis was performed using the RT² First Strand Kit (Qiagen) with 1000 ng of total RNA per reaction. Gene expression profiling was carried out using RT² Profiler PCR Array plates (Qiagen) specific for Human Endothelial Cell Biology (PAHS-015Z) and Human Hematopoiesis (PAHS-054Z), and qPCR reactions were set up using RT² SYBR Green qPCR Mastermix (Qiagen). Ct values were analyzed using Qiagen’s RT² Profiler PCR Array Data Analysis Web Portal (https://geneglobe.qiagen.com/us/analyze), using the ΔΔCt method. Gene expression was normalized to the geometric mean of housekeeping genes provided on the array. Genes with a fold change ≥1.5 and p-value <0.05 were considered significantly differentially expressed.

### Western blot analysis

Cells were lysed in RIPA buffer (Thermo Fisher Scientific, Cat# 89900) containing protease inhibitors. Protein levels were quantified using the BCA assay (Thermo Fisher Scientific, Cat# 23227), and equal amounts (20-40 µg) were separated by 4-20% SDS-PAGE gel and transferred to 0.2 µm nitrocellulose membranes using the Trans-Blot Turbo Transfer System (Bio-Rad, Cat# 1704150). Membranes were blocked in 5% milk/TBS-T, incubated overnight at 4°C with primary antibodies (Supplementary Table S2) in 5% BSA/TBS-T, and then with IRDye-conjugated secondary antibodies (LI-COR). Signals were detected on the Odyssey CLx Imaging System, and band intensities were quantified with Image J software.

### Endothelial tube formation assay

Tube formation ability of HPAECs was assessed using the Cultrex® Basement Membrane Extract (BME, R&D Systems, Cat# 3432-005-01). A clear, flat-bottom 96-well tissue culture plate (pre-cooled on ice) was coated with 70 µl of cold BME per well and incubated at 37°C for 30 m to allow gel polymerization. After gelation, 10,000 to 15,000 HPAECs in 100 µl of complete growth medium were seeded into each well. Cells were incubated at 37°C in a 5% CO₂ humidified incubator for 16 h. Tube formation was monitored using an inverted phase contrast microscope (4x and 10x objectives), and representative images were captured at indicated time points. Quantification of tube formation parameters was performed using ImageJ software with the Angiogenesis Analyzer plugin.

### Endothelial cell migration assay

Endothelial cell migration was assessed using the Ibidi 2-Well Culture Inserts (Ibidi, Cat# 80209) placed in a 12-well tissue culture plate. HPAECs were seeded into each well of the insert at a density of 90,000 cells in 90 µl of complete growth medium. Cells were incubated overnight at 37°C in a humidified incubator with 5% CO₂ to allow the attachment. Following incubation, the insert was carefully removed using sterile forceps to create a defined cell-free gap. Wells were gently washed with pre-warmed PBS to remove detached cells and replaced with fresh medium. Migration was monitored by capturing images with an inverted phase contrast microscope with 4x magnification at defined time points (e.g., 0, 6, and 12 h). Gap closure was expressed as the percentage of the initial wound area covered by migrating cells over time using ImageJ software.

### MTS proliferation assay

HPAECs were seeded in clear TC-treated 96-well plates at a density of 4,000 cells/well in complete EGM-2 media and allowed to adhere and proliferate, and cell density was assessed at indicated days using the CellTiter 96® AQueous One Solution Cell Proliferation Assay Reagent (Promega, Cat# G3582), according to the manufacturer’s instructions. Briefly, 20 µl of MTS reagent was added directly to each well containing 100 µl of medium, mixed well and incubated for 2 h at 37°C in in dark, in a humidified atmosphere with 5% CO₂. Absorbance was measured at 490 nm using a microplate reader. Background absorbance (wells containing medium and MTS reagent but no cells) was subtracted from all readings for calculation.

### Caspase 3/7 activity assay for apoptosis

Apoptosis was assessed using the Caspase-Glo® 3/7 Assay kit (Promega, Cat# G8091) according to the manufacturer’s instructions. Briefly, HPAECs were seeded in white, opaque-walled 96-well plates at a density of 10,000 cells per well in 100 µl of media and allowed to adhere overnight. To induce apoptosis, cells were treated with human TNF-α (PeproTech, Cat# 300-01A) at a final concentration of 10 ng/ml for 16 h. Following incubation, an equal volume of Caspase-Glo® 3/7 reagent was added directly to each well. Plates were gently mixed and incubated for 30 m at room temperature in the dark. Luminescence readings were recorded using a microplate reader. Background signal from wells containing medium and reagent but no cells was subtracted from sample readings. All conditions were tested in triplicates, and data were normalized to untreated controls.

### Small molecule RUNX1 inhibitors prevent the development of SuHx-PH in SOX17enhKO mice

As a model of PH we used transgenic mice with deletion of SOX17 signal 1 (SOX17 enhancer knock out, SOX17enhKO), which are designed to mimic the common risk variants associated with human PAH and develop more severe PH when exposed to hypoxia ^13^. Following exposure to mild hypoxia (12% O2) and low dose Sugen 5416 (5 mg/kg) these mice develop severe PH which causes no PH in wild-type mice. To determine if small molecule RUNX1 inhibitors are able to abrogate the susceptibility of SOX17enhKO mice in SuHx-PH, we subcutaneously injected Ro5-3335 (20 mg/kg, MedChemExpress, Cat# HY-108470), Ro24-7429 (40 mg/kg, MedChemExpress, Cat# HY-19149) or DMSO vehicle control every other day for 6 times starting 1 week after exposure to low dose Sugen/hypoxia (Figure 9A). Endpoints were measured at 3 weeks. Development of PH was determined by measurement of right ventricular systolic pressure (RVSP), right ventricular (RV) hypertrophy [RV to left ventricle + septum wet weight (RV/LV+S) ratio] and pulmonary arteriole remodeling. To measure RVSP, mice were anesthetized via continuous inhalation of 1-3% isoflurane. An 1 cm incision was made carefully in the neck skin and the right internal jugular vein was isolated by using eye-dressing forceps. A tiny incision was made at the proximal end (towards the heart) of the jugular vein and a pressure transducer (Cat # PVR-1030, Millar) was inserted and slowly advanced into the right ventricle. RVSP measurements were collected and analyzed using the PowerLab 4/35 four-channel data acquisition system with LabChart Pro software (Millar). Mice were then sacrificed by exsanguination, heart and lung were removed and fixed in 10% formalin solution.

Immunohistochemical staining (IHC) of mouse α-smooth muscle actin was carried out on paraffin embedded lung sections by using a rabbit polyclonal antibody (Abcam ab5694, 1:200). IHC images were captured through full slide scanning. NIH ImageJ software was then used to calculate muscularization indices, which were defined as [(outer area-inner area)/outer area] ratio, for pulmonary arterioles (≤ 50 µm). All animal experiments were conducted in accordance with institutional guidelines and approved by the Institutional Animal Care and Use Committee (IACUC) of Brown University Health.

### Endothelial specific deletion of RUNX1 prevents the development of SuHx-PH in SOX17enhKO mice

In order to inducibly delete RUNX1 in adult endothelium, we crossbred Cdh5(PAC)-CreERT2 mice ^28^ (Taconic Cat# 13073, Rensselaer, NY) with Runx1(flox/flox) mice (Jackson Laboratory Cat# 008772, Bar Harbor, ME) and generated Cdh5-CreERT2;Runx1(flox/flox) mice in our lab ^21^. Mice with correct genotype were each treated with 2 mg of tamoxifen daily via intraperitoneal injection for 5 days followed by 1 week of incubation. Lung ECs from the mice treated with tamoxifen were isolated via flow cytometry cell sorting ^29^, and the loss of RUNX1 in ECs was verified per qRT-PCR with the following primers: forward 5’- CCT CCT TGA ACC ACT CCA CT - 3’, and reverse 5’- CTG GAT CTG CCT GGC ATC -3’. GAPDH expression was assessed per qRT-PCR as an internal reference for quantification with the following primers: forward 5’ - AAA AGC AAC TCC CAC TCT TC - 3’ and reverse 5’ - CCT GTT GCT GTA GCC GTA TT - 3’. All PCR primers were synthesized by Integrated DNA Technologies (Coralville, IA).). As a control for endothelial specificity of RUNX1 deletion, we confirmed RUNX1 gene expression in CD14+ monocytes isolated from bone marrow (BM) of these mice following tamoxifen induction ^21^. An anti-mouse CD14 antibody conjugated with APC was purchased from BioLegend and BM CD14+ cells were sorted on a BD Influx flow cytometry cell sorter (BD Biosciences). We then crossbred the Cdh5-CreERT2;Runx1(flox/flox) mice with SOX17enhKO mice to generate triple transgenic Cdh5-CreERT2;Runx1(flox/flox);SOX17enhKO mice (named as SRV mice in this study: S as in SOX17, R as in RUNX1 and V as in VE-Cadherin). Prior to SuHx-PH induction in the SRV mice, tamoxifen or vehicle corn oil was given daily via intraperitoneal injection for 5 days followed by 1 week of incubation. The mice were then subjected to 3 weeks of mild hypoxia (12% O_2_) and weekly injection of low dose Sugen 5416 (5 mg/kg). Development of PH in mice was determined as described above.

### Small molecule RUNX1 inhibitors Ro5-3335 and Ro24-7429 reversed SuHx-PH in rats

The SuHx-PH model in rats was carried out as published previously by us ^21, 30^. Briefly, male Sprague-Dawley rats (Charles River Laboratory, Wilmington, MA) weighing 125-150 g were injected subcutaneously with 25 mg/kg of the VEGF receptor 2 antagonist Sugen 5416 (SU5416, Bio-Techne, Minneapolis, MN) and placed in hypoxic chambers (10.5% O2) for 3 weeks followed by 2 weeks of normoxic recovery. Normoxia (Nx) control rats were injected with an equal volume of dimethyl sulfoxide (DMSO) vehicle, kept under normoxic conditions, and then injected with an equal volume of DMSO at the same time points when SuHx-PH rats were given RUNX1 inhibitor. In the disease reversal protocol, rats were given 20 mg/kg of Ro5-3335, or 40 mg/kg of Ro24-7429 in 100 µl of DMSO or 100 µl vehicle alone by subcutaneous injection every other day for 6 times after the completion of SuHx treatment, and the development of PH was assessed 2 weeks after removal from hypoxia (Supplementary Figure S3A).

### Statistical analysis

All statistical analyses were performed using GraphPad Prism version 11 (GraphPad Software, San Diego, CA). Data are presented as mean ± Standard Error of the Mean (SEM). Comparisons between two groups were conducted using unpaired t-tests. One-way ANOVA was employed for comparisons among multiple groups. A p-value of less than 0.05 was considered statistically significant.

## RESULTS

### SOX17 deficiency in HPAECs leads to loss of endothelial identity and endothelial dysfunction

CRISPR/Cas9 knockout of SOX17 (SOX17 KO) in human pulmonary artery endothelial cells (HPAECs) resulted in near complete deletion of SOX17 (Figure 1A). Compared to HPAECs treated with control vector, SOX17 KO HPAECs exhibited decreased expression of numerous endothelial cell markers including PECAM (CD31), VE-cadherin (CD144), ICAM-1, von Willebrand factor (vWF), and KDR (VEGFR2) (Figures 1A, B and C). Other genes involved in endothelial cell growth and function such as VEGFA, TGFB1, endostatin (COL18A), and ENG were also down regulated, as were END1 and EDNRA which encode endothelin-1 and its receptor endothelin receptor type A, respectively and CFLAR which encodes c-FLIP major inhibitor of caspase-8 and apoptosis. SOX17 KO also had marked effects on genes involved in vascular coagulation with increased expression of PLAT which encodes for tissue-type plasminogen activator (tPA) and down regulation of SERPINE1 which encodes plasminogen activator inhibitor-1 (PA-1) which is the primary inhibitor of tPA. Differentially expressed genes with the greatest difference from control included the growth factors IL-1β and FGF1 which play major roles in endothelial cell survival and angiogenesis.

**Figure 1.**
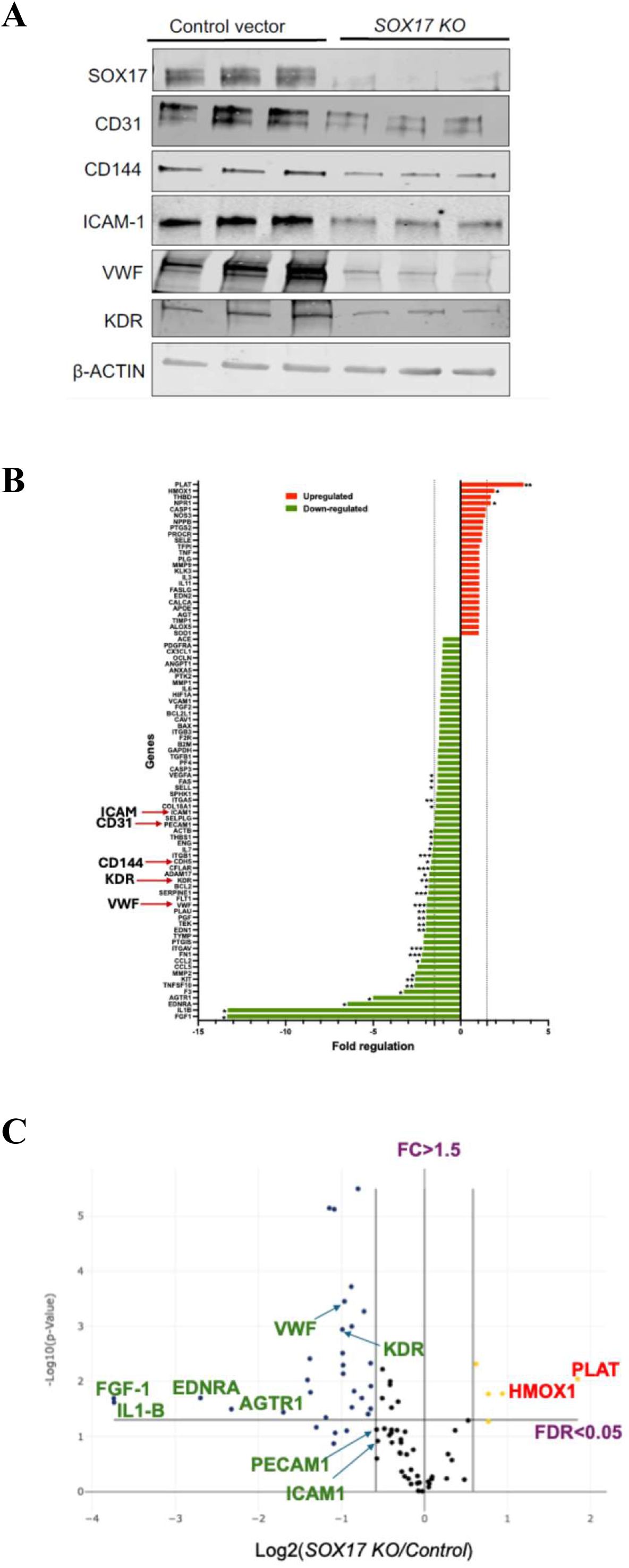
SOX17 deficiency in HPAECs leads to loss of endothelial cell markers. HPAECs subjected to CRISPR/Cas9-mediated SOX17 knockout (SOX17 KO) were utilized to define SOX17-dependent changes in endothelial cell biology. (A) Representative western blot analysis of key endothelial cell markers in control cells versus two independent SOX17 KO clones (SOX17 KO-1 and KO-2). (B and C) Human Endothelial Cell Biology RT²-PCR Array analysis comparing SOX17 KO HPAECs with control vector transduced cells. Bar graphs and volcano plots highlight significantly dysregulated genes (fold change > 1.5, FDR < 0.05).

Compared to control cells, SOX17 KO HPAECs exhibited loss of cobblestone morphology, and a more spindle appearance with increased length/width ratio (Figure 2A). Tube formation assays revealed impaired angiogenic network assembly in SOX17 KO HPAECs as indicated by reduced number of cell junctions, reduced number of meshes, smaller mesh area, shorter tube length and fewer nodes (Figure 2B). SOX17 KO in HPAECs also led to enhanced migration (Figure 2C), increased proliferation (Figure 2D), and decreased apoptosis at baseline and in response to TNF-α compared to control cells (Figures 2E).

**Figure 2.**
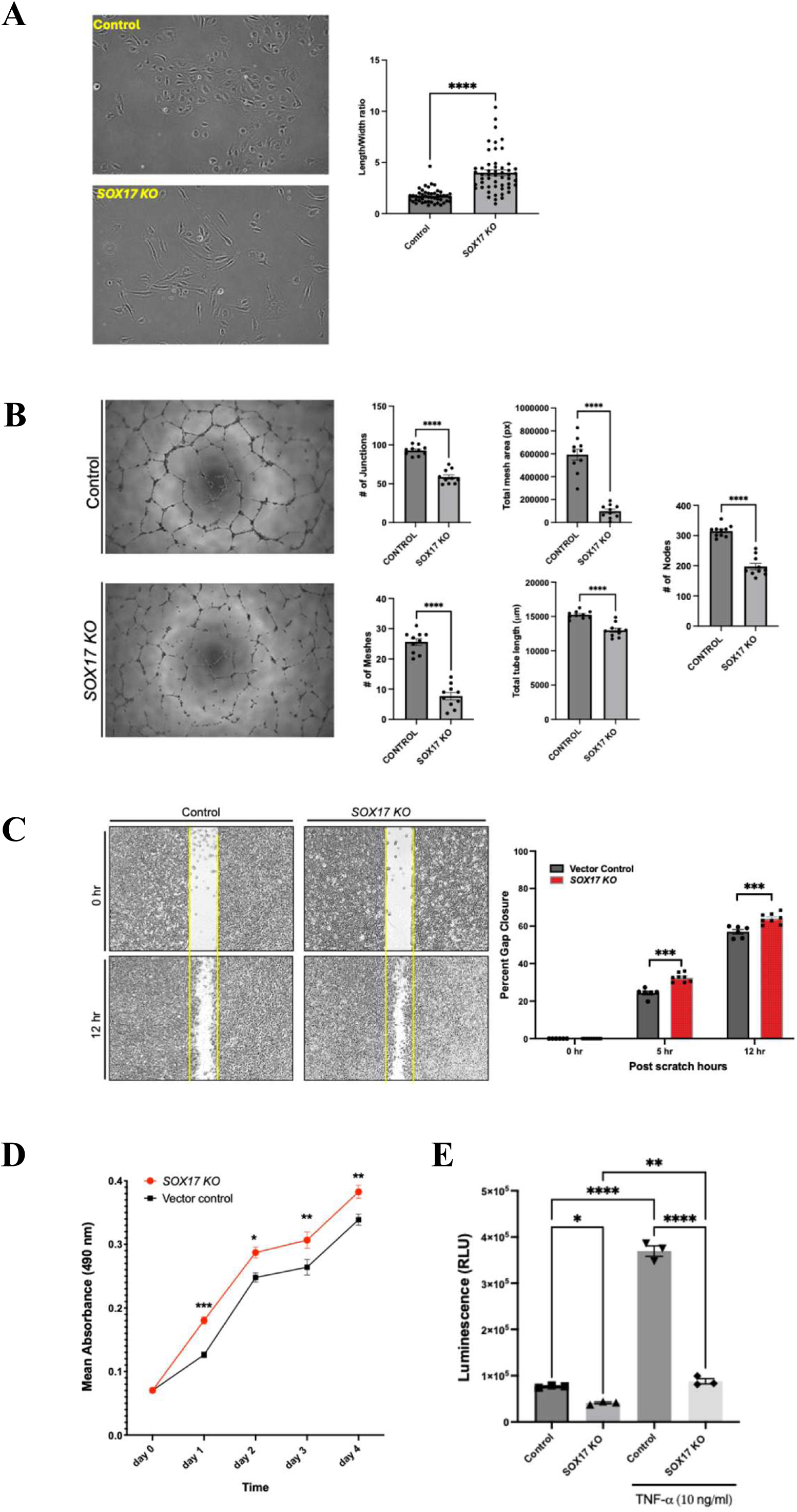
SOX17 deficiency in HPAECs causes endothelial dysfunction. (A) Morphometric analysis (length-to-width ratio) demonstrated marked alterations in cell shape and loss of the characteristic cobblestone morphology in SOX17 KO HPAECs. (B) Tube formation assays revealed impaired angiogenic network formation in SOX17-deficient cells. (C) Migration assays showed accelerated gap closure in SOX17 KO PAECs, consistent with enhanced motility. (D) MTS assays demonstrated increased proliferative capacity. (E) Reduced Caspase-3/7 activity following TNF-α treatment indicated decreased apoptosis in SOX17 KO HPAECs. All image-based analyses (A-C) were quantified using ImageJ software. Statistical significance for (D and E) was determined by two-way ANOVA with multiple comparisons, while all other comparisons were performed using unpaired two-tailed t-tests. *p < 0.05, **p < 0.01, ***p < 0.001, ****p < 0.0001.

### RUNX1 overexpression in HPAECs leads to loss of endothelial identity and endothelial dysfunction

Transduction of HPAECs with lentivirus harboring the RUNX1 overexpression construct significantly increased RUNX1 gene expression and protein levels compared to control lentivirus (Figure 3A). Similar to SOX17 KO, overexpression of RUNX1 in HPAECs resulted in significant downregulation of numerous genes associated with endothelial identity and endothelial function including vWF, PECAM, VE-cadherin, and ICAM-1 (Figure 3A, B and C). Other genes involved in endothelial cell growth and function downregulated by SOX17 KO were also similarly downregulated by RUNX1 overexpression including VEGFA, TGFB1, endostatin (COL18A), ENG, END1, EDNRA, SERPINE-1, and CFLAR (Figure 3B and C). PLAT which was upregulated by SOX17 KO was also upregulated by RUNX1 overexpression. Except for IL-1β, which was markedly downregulated by SOX17 KO but upregulated by RUNX1 overexpression, the great majority of endothelial specific genes were regulated in the same direction by SOX17 KO and RUNX1 overexpression. Overexpression of RUNX1 in HPAECs resulted in the loss of endothelial cell cobblestone morphology as demonstrated by spindle cell shape with increased length to width ratio (Figure 4A). RUNX1 overexpression also caused impaired tube formation (Figure 4B), increased cell migration (Figure 4C), cell proliferation (Figure 4D), and decreased apoptosis (Figure 4E).

**Figure 3.**
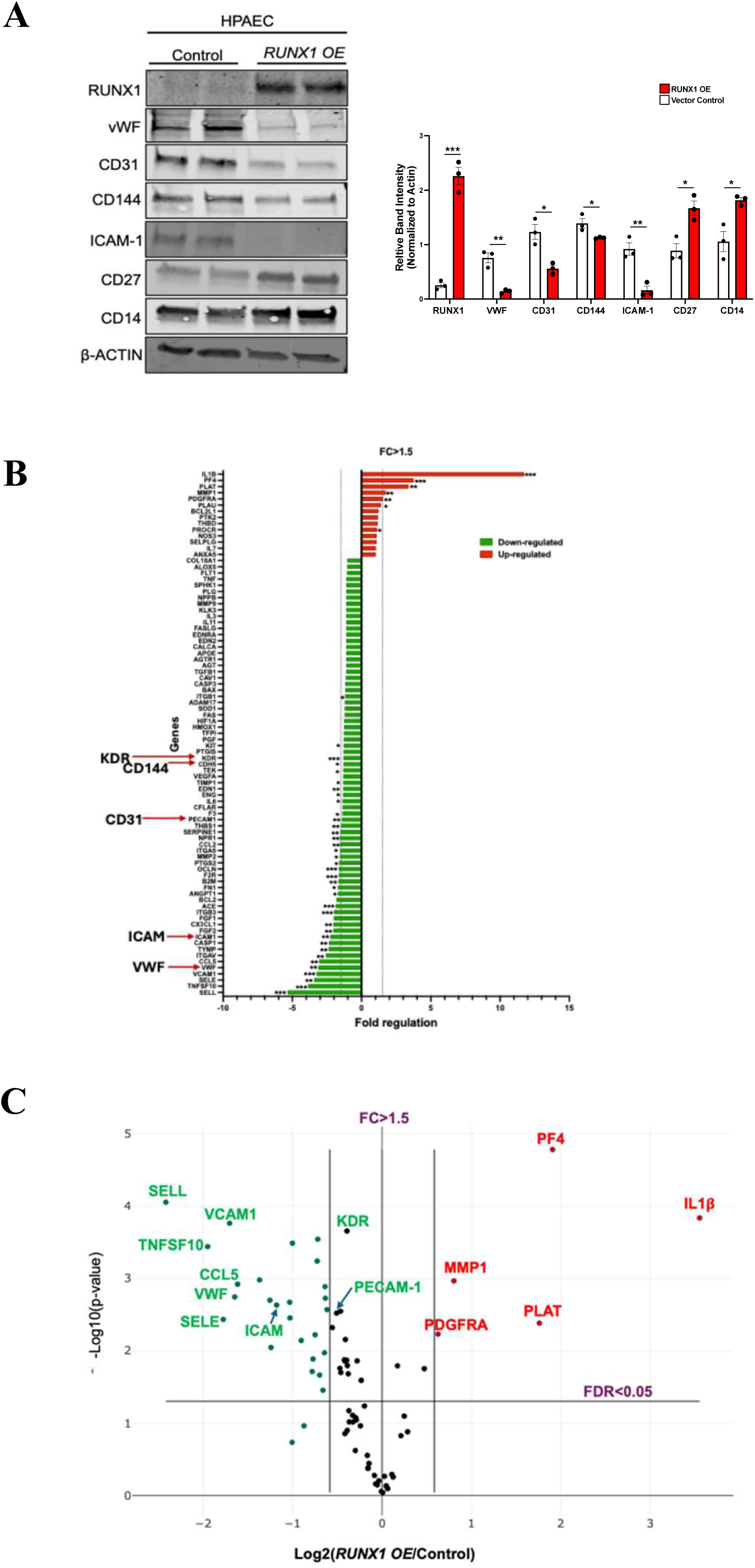
RUNX1 overexpression drives a transcriptional shift away from endothelial identity in HPAECs. (A) Representative Western blot analyses of endothelial and hematopoietic markers in control versus RUNX1 overexpression HPAECs, with quantification of relative band intensities normalized to loading controls. (B and C) Human Endothelial Cell Biology RT²-PCR Array analyses in HPAECs overexpressing RUNX1 compared to control vector transduced cells. Gene expression values were normalized to housekeeping genes and represented as fold change relative to controls. Bar graphs and Volcano plots highlight differentially expressed genes meeting the significance criteria of fold change > 1.5 and FDR < 0.05. *p < 0.05, **p < 0.01, ***p < 0.001 indicate statistical significance.

**Figure 4.**
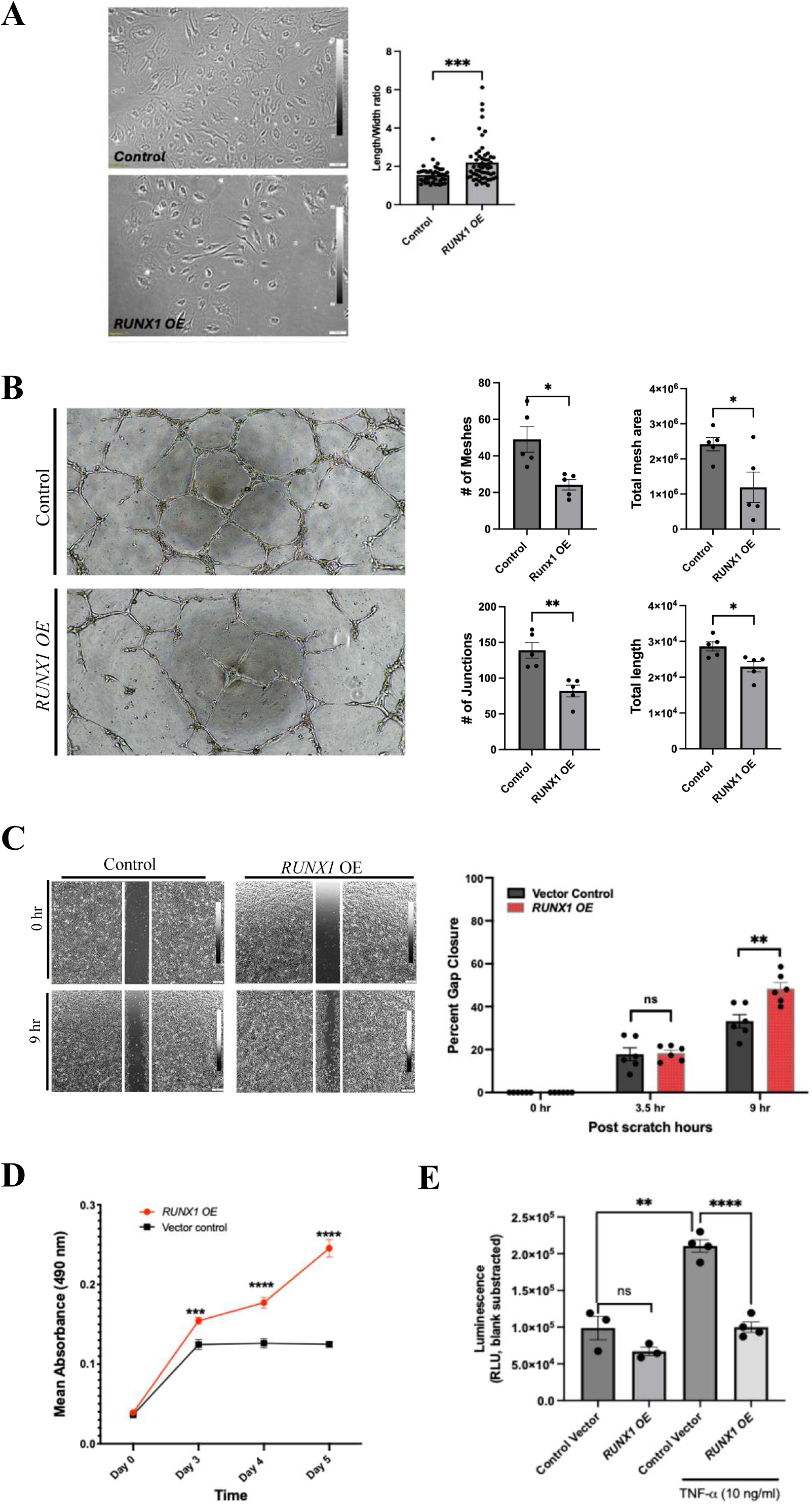
RUNX1 overexpression in HPAECs causes endothelial dysfunction. HPAECs transduced with RUNX1 overexpressing (RUNX1 OE) or control lentivirus were subjected to quantitative assays to evaluate RUNX1-driven phenotypic and functional changes. (A) Morphometric assessment (length/width ratio) demonstrated pronounced alterations in cell morphology in RUNX1 OE cells. (B) Tube formation assays demonstrated impaired angiogenic network assembly (C) Migration assays showed increased gap closure over time in RUNX1 OE cells. (D) MTS proliferation assays indicated enhanced growth in RUNX1 OE cells. (E) Reduced Caspase-3/7 activity following TNF-α treatment demonstrated decreased apoptosis in RUNX1 OE cells. All image-based analyses (A-C) were quantified using ImageJ software. Statistical significance for (D and E) was determined by two-way ANOVA with multiple comparisons, while all other comparisons were performed using unpaired two-tailed t-tests. *p < 0.05, **p < 0.01, ***p < 0.001, ****p < 0.0001.

In addition, RUNX1 overexpression promoted a shift away from endothelial identity by activating a hematopoietic transcriptional program in HPAECs, as reflected by increased expression of multiple hematopoietic marker genes (Supplementary Figure S1). Specifically, HPAECs with RUNX1 overexpression led to significant upregulation of hematopoietic gene expression including lymphocyte markers CD4 and CD27, megakaryocyte and platelet marker PF4, and monocyte marker CD14, as well as significant downregulation of stem and progenitor cell marker CD34 (Figure 3A and Supplementary Figure S1).

### ECs derived from PAH patients with SOX17 mutations have reduced expression of endothelial cell markers

Induced pluripotent stem cells (iPSCs) were generated from lymphoblastoid cell lines (LCLs) obtained from patients with SOX17 mutations and commercially available LCLs from healthy donors. Characterization of the iPSCs was performed to demonstrate: 1) expression of pluripotency markers c-MYC, KLF4, LIN28A, OCT4A, SOX2, and NANOG in Western blots (Supplementary Figure S2A), 2) immunocytochemical staining of pluripotency markers OCT4 (red), SSEA4 (green), SOX2 (green), and TRA-1-60 (red), with DAPI counterstaining of nuclei (blue) (Supplementary Figure S2B), 3) trilineage differentiation potential of healthy control and PAH LCL-derived iPSCs by immunocytochemistry, where differentiated cells were stained for ectoderm lineage marker Nestin (green), mesoderm lineage marker Brachyury (red), and endoderm lineage marker FOXA2 (red), with DAPI counterstaining of nuclei (blue) (Supplementary Figure S2C), 4) normal chromosomal integrity in karyotype analysis (Supplementary Figure S2D). Endothelial cells were derived from the respective iPSCs (Figure 5A). Endothelial cells derived from iPSCs from PAH patients with SOX17 mutation exhibited altered morphology (Figure 5B) and reduced expression of a number of endothelial markers including ICAM, vWF, KDR (VEGFR2), CD144 (VE-cadherin) and CD31 (PECAM) (Figure 5C and D).

**Figure 5.**
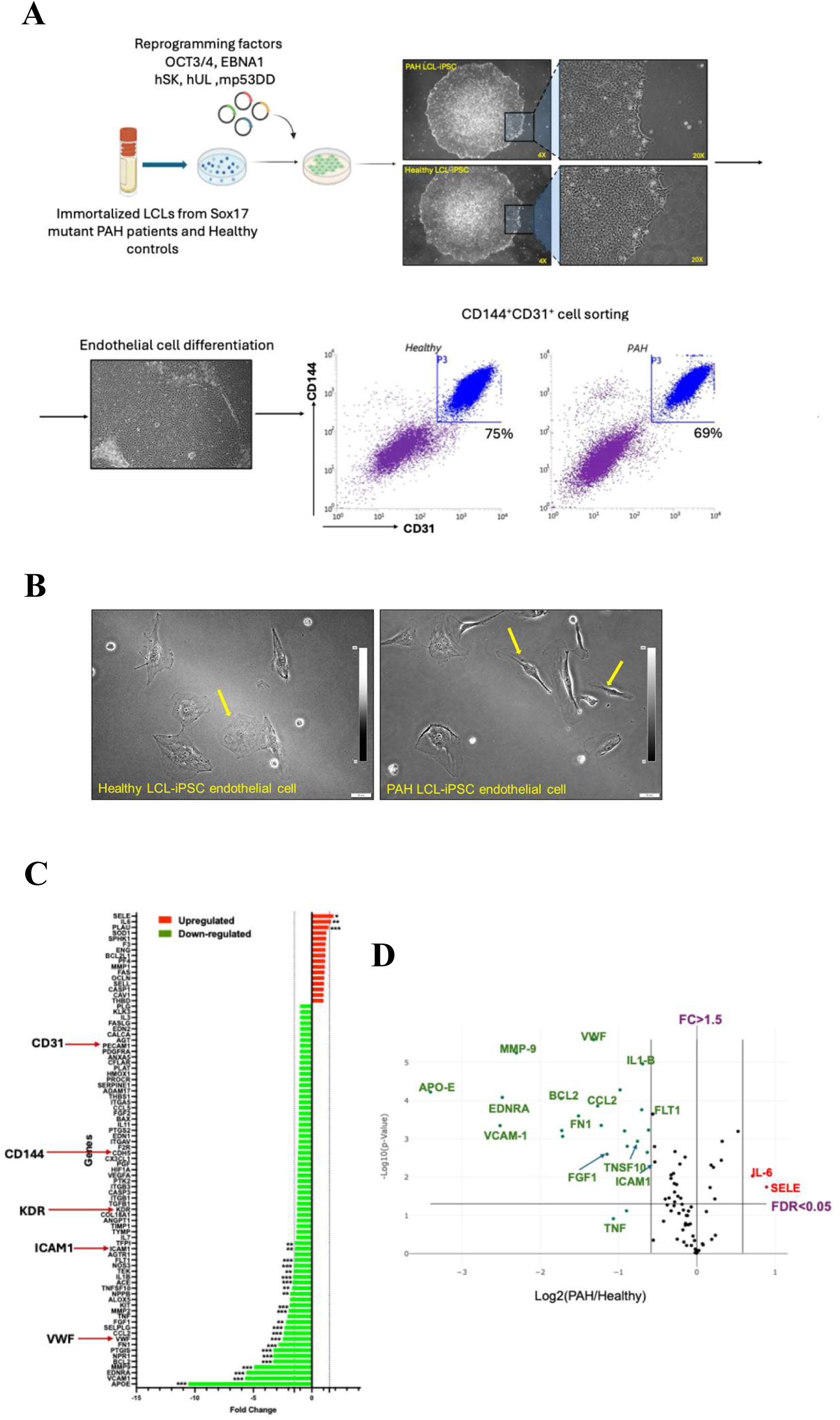
ECs derived from PAH patients with SOX17 mutations have reduced expression of endothelial cell markers. (A) Schematic overview of iPSC generation from lymphoblastoid cell lines (LCLs) of healthy controls and SOX17-mutant PAH patients, followed by endothelial differentiation and FACS sorting. (B) Morphological appearance of sorted CD31+CD144+ endothelial cells (scale bar, 50 µm). (C and D) Human Endothelial Cell Biology RT²-PCR Array profiling of SOX17 mutant iPSC-ECs compared with healthy controls. Volcano plots and bar graphs depict differentially expressed genes meeting the criteria of fold change > 1.5 and FDR < 0.05. *p < 0.05, **p < 0.01, ***p < 0.001, ****p < 0.0001 denote statistical significance.

### RUNX1 expression is increased in ECs with impaired SOX17 expression

RUNX1 protein level was significantly higher in SOX17 KO HPAECs compared to control HPAECs (Figure 6A). As shown earlier in Figure 1, SOX17 KO HPAECs had significantly reduced levels of endothelial markers vWF, ICAM, and VE-cadherin (CD144). This effect was partially reversed by treatment with either the RUNX1 inhibitor Ro5-3335 (Figure 6B) or by siRNA knockdown of RUNX1 (Figures 6C).

**Figure 6.**
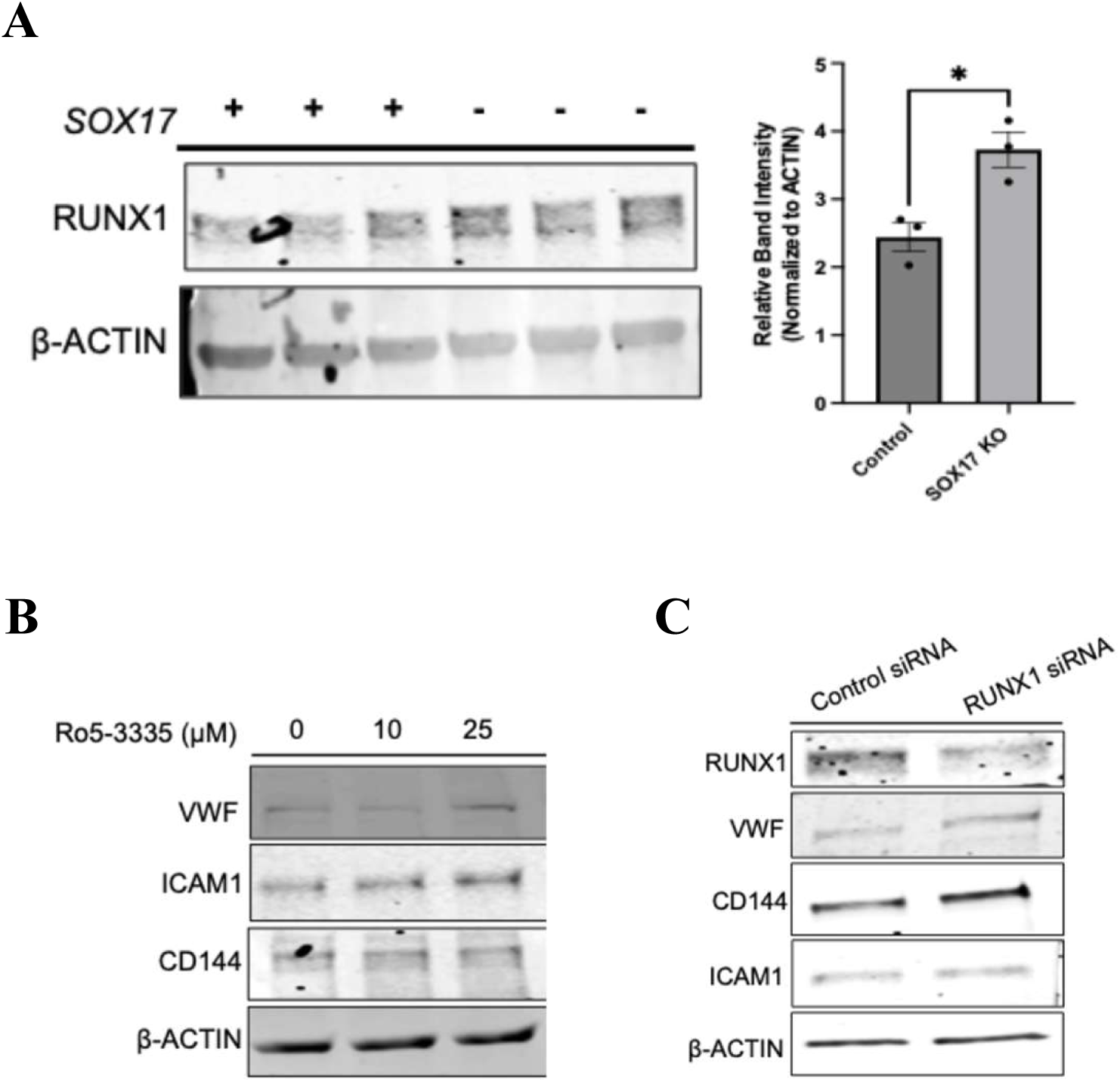
Upregulation of RUNX1 expression in SOX17 KO HPAECs. Western blot analyses show (A) increased RUNX1 protein levels in SOX17 KO HPAECs, with quantification of relative band intensity normalized to β-ACTIN. (B and C) Treatment of SOX17 KO HPAECs with RUNX1 inhibitor Ro5-3335 at 0, 10 or 25 µM (B) or RUNX1-targeting siRNA (C) leads to partial restoration of endothelial marker expression in the absence of SOX17. *p < 0.05.

Furthermore, both mRNA (Figure 7A) and protein (Figure 7B) levels of RUNX1 were greater in iPSC-ECs derived from PAH patients with SOX17 mutations compared to iPSC-ECs from healthy controls. Similar to HPAECs with SOX17 KO, iPSC-ECs from PAH patients with SOX17 mutations had decreased expression of endothelial markers such as vWF, ICAM-1 and VCAM (Figure 7B). Treatment of iPSC-ECs from PAH patients with SOX17 mutations with either the RUNX1 inhibitor Ro5-3335 (Figure 7C) or by RUNX1 siRNA (Figure 7D) increased expression of EC markers.

**Figure 7.**
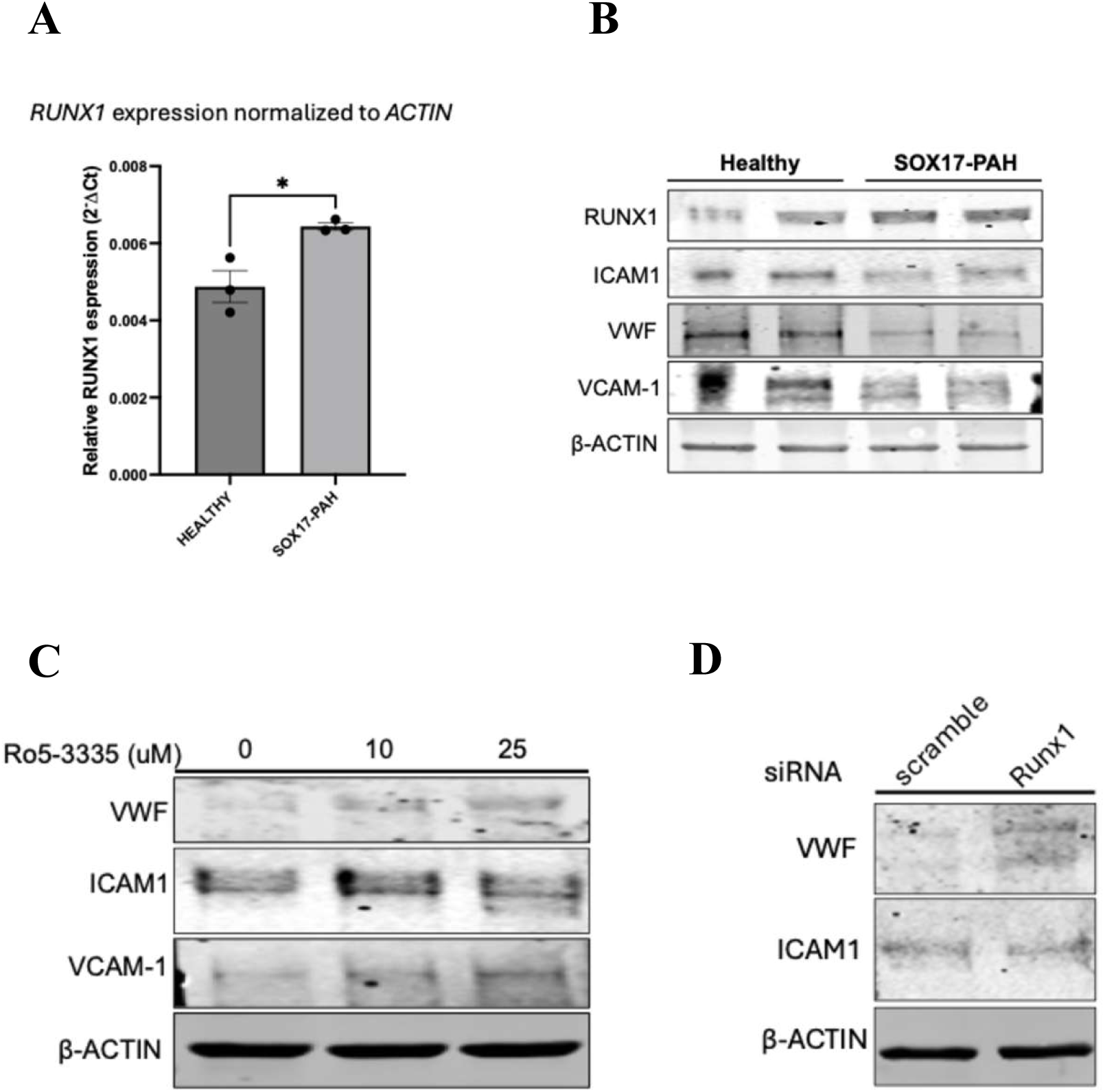
Upregulation of RUNX1 expression in iPSC-ECs derived from SOX17 mutant PAH patients. (A and B) RT-PCR analyses (A) and Western blot analyses (B) demonstrate increased RUNX1 expression and reduced endothelial marker levels in iPSC-ECs derived from SOX17 mutant patients compared with healthy controls. (C) Western blots shows in a dose-dependent manner RUNX1 inhibitor Ro5-3335 partially restored endothelial marker expression in iPSC-ECs derived from SOX17 mutant PAH patients. (D) Western blots shows RUNX1 targeting siRNA downregulation partially restored endothelial marker expression in iPSC-ECs derived from SOX17 mutant PAH patients. *p < 0.05, **p < 0.01, ***p < 0.001, ****p < 0.0001 denote statistical significance.

### RUNX1 expression is increased in SOX17enhKO mice with SuHx-PH

In a prior study, we showed that SOX17enhKO mice, designed to mimic common SOX17 variants that are associated with increased risk of PAH in humans, develop more severe pulmonary hypertension in response to chronic hypoxia than wild-type mice, and are more susceptible to the development of SuHx-PH ^13^. These mice develop PH in response to a reduced dose of Sugen 5416 (5 mg/kg) and 3 weeks of mild hypoxia (12.5% oxygen) that is just as severe as that induced by 20 mg/kg Sugen 5416 and 8.5% oxygen in wild-type mice. This reduced dose of Sugen and less severe hypoxia causes no PH in wild-type mice.

To determine if the increased susceptibility of SOX17enhKO mice is mediated by RUNX1, we measured RUNX1 expression in the bone marrow and lung of SOX17enhKO and wild-type mice in the absence and presence of low dose Sugen/hypoxia. In mice that were not exposed to low dose Sugen/hypoxia, no difference in RUNX1 expression in bone marrow was seen between SOX17enhKO and wild-type mice (Figure 8A). Treatment with mild Sugen/hypoxia decreased RUNX1 expression in the bone marrow of wild-type mice, but increased RUNX1 in bone marrow of SOX17enhKO mice. Similarly, RUNX1 expression in the lung did not differ between SOX17enhKO and wild-type mice under normoxic conditions and treatment with low dose Sugen/hypoxia increased RUNX1 expression in the lungs of SOX17enhKO mice but not in wild-type mice (Figure 8B).

**Figure 8.**
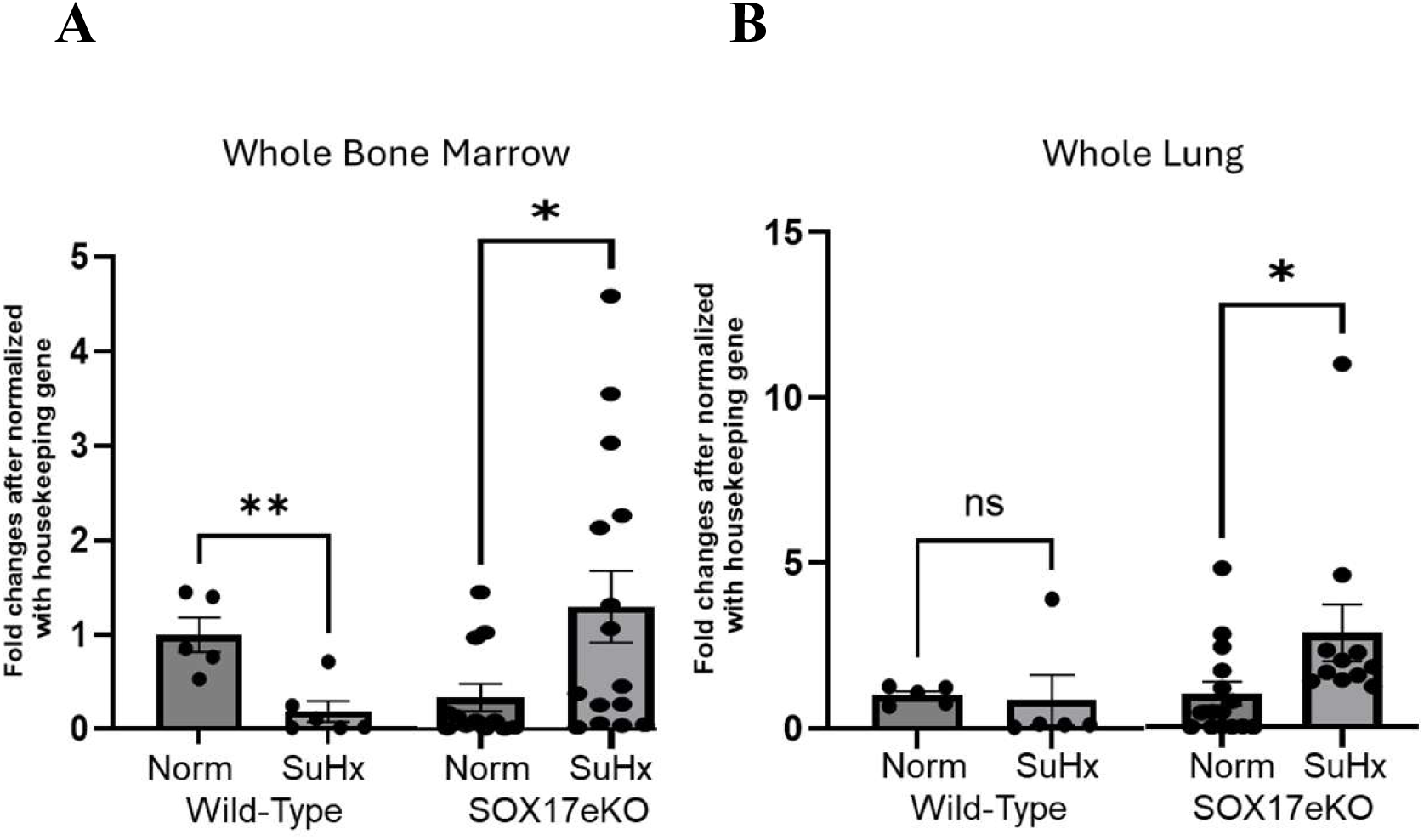
RUNX1 expression is increased in SOX17enhKO mice under SuHx. RUNX1 expression in whole bone marrow (A) and whole lung (B) in SOX17enhKO and WT mice after 3 weeks of normoxia (Norm) or Sugen/hypoxia (SuHx). N = 5-15 mice per group, * P < 0.05, ** P < 0.01, ns: not significant.

### RUNX1 inhibition decreases the susceptibility of SOX17enhKO to SuHx-PH

To determine if the increased susceptibility to mild Sugen/hypoxia exhibited by SOX17enhKO mice can be mitigated by inhibition of RUNX1, we treated SOX17enhKO and wild-type mice with the RUNX1 inhibitors Ro5-3335 or Ro24-7429. Mice were exposed to 12.5% oxygen and injected with 5 mg/kg Sugen 5416 once weekly for 3 weeks or kept in normoxia and injected with Sugen vehicle (DMSO) alone. One week into SuHx treatment, mice were treated with the RUNX1 inhibitors or vehicle alone by subcutaneous injection every other day for 6 times (Figure 9A). Mild Sugen/hypoxia caused severe PH in SOX17ehKO mice as evidenced by increases in RVSP and RV/(LV+S) but not in wild-type mice (Figure 9B and C). Treatment with Ro5-3335 or with an alternative RUNX1 inhibitor Ro24-7429, completely prevented the Sugen/hypoxia-induced increase in RVSP and significantly reduced the increase in RV/(LV+S) (Figure 9B and C), suggesting that the increased susceptibility to Sugen/hypoxia-PH in SOX17enhKO mice can be rescued by inhibiting the effect of increased RUNX1 expression.

**Figure 9.**
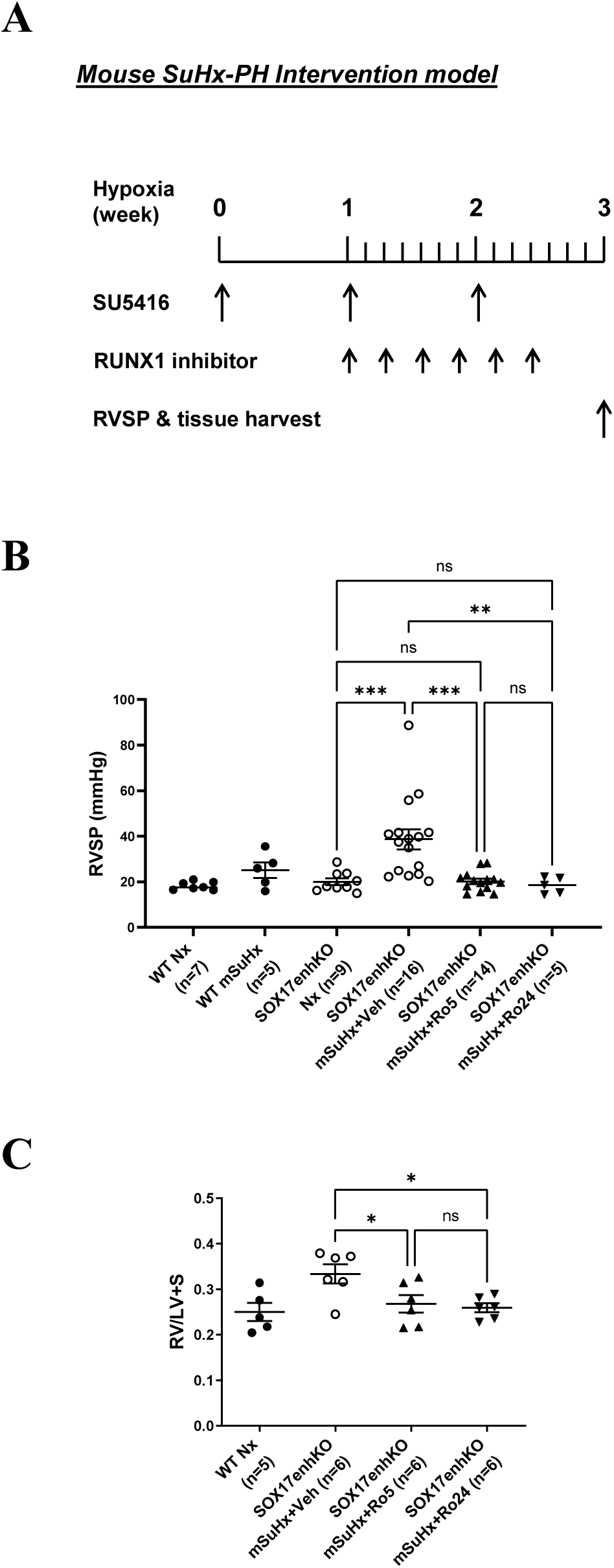
Small molecule RUNX1 inhibition rescued the susceptibility of SOX17enhKO mice in developing SuHx-PH. (A) Experimental protocol for intervention of SuHx-PH in SOX17enhKO mice shows administration of the RUNX1 inhibitor Ro5-3335 or Ro24-7429 every other day for 6 times 1 week after the beginning of mild SuHx treatment. (B and C) RVSP (B) and RV/LV+S ratio (C) were measured at the end of week 3. Data in (B) and (C) are mean ± SEM. * P < 0.05, ** P < 0.01, *** P < 0.001, ns: not significant.

### Endothelial specific deletion of RUNX1 abrogated the susceptibility of SOX17enhKO mice in developing SuHx-PH

To ensure that the modulatory effect of Ro5-3335 or Ro24-7429 on Sugen/hypoxia-PH is mediated by their inhibitory effect on RUNX1 and not by off target inhibition of other transcription factors or proteins, we performed additional experiments in Cdh5-CreERT2;Runx1(flox/flox);SOX17enhKO triple transgenic SRV mice, which provided inducible endothelial specific knockout of RUNX1 in the SOX17ehKO mice. Mild Sugen/hypoxia had no effect on wild-type (WT) mice but caused significant increases in RVSP and (RV/LV+S) in triple transgenic SRV mice that were not given tamoxifen (Figures 10). Endothelial deletion of RUNX1 by administration of tamoxifen prevented the Sugen/hypoxia induced increase in RV pressure (Figure 10).

**Figure 10.**
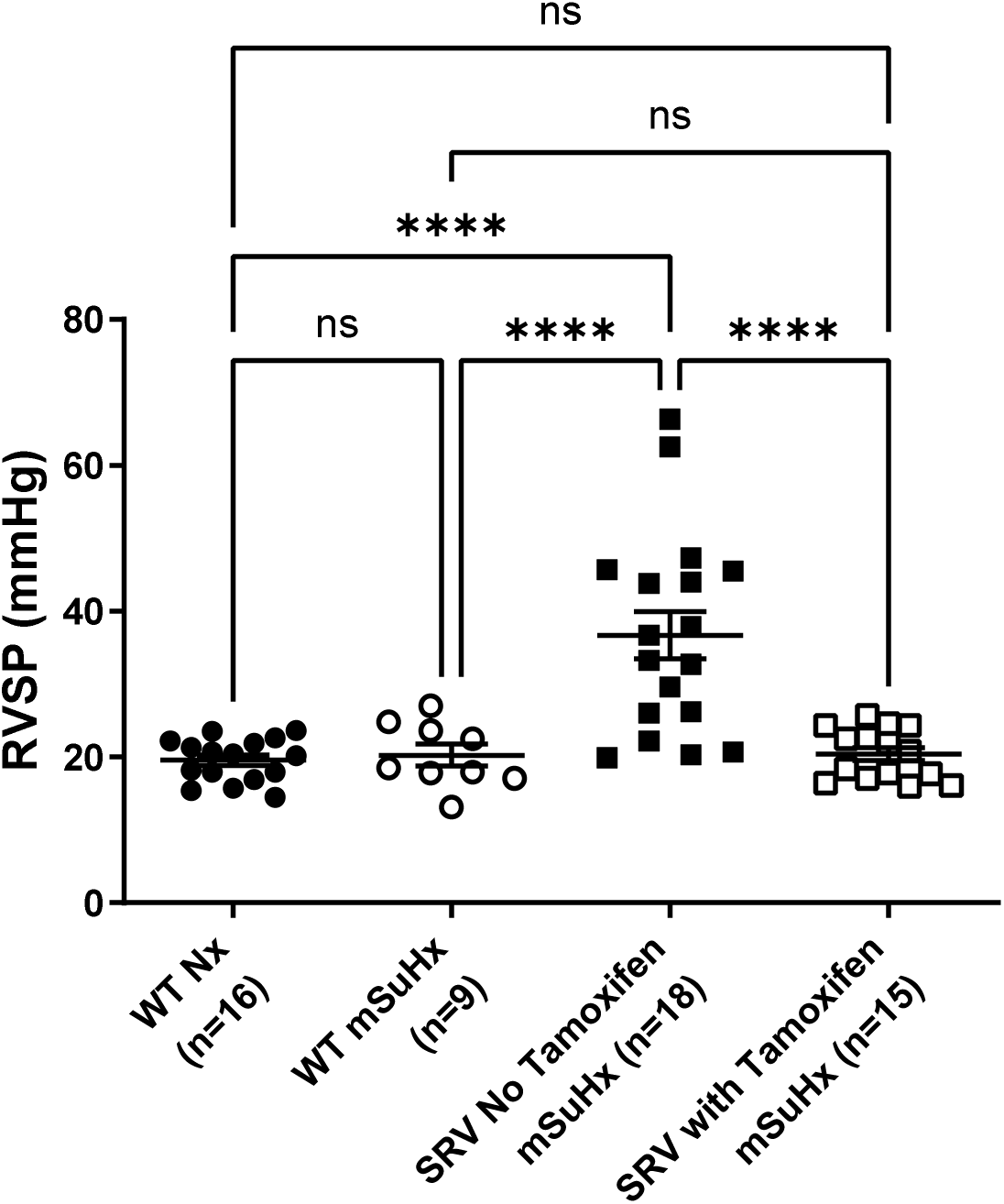
Endothelial specific deletion of RUNX1 abrogated the susceptibility of SOX17enhKO mice in developing SuHx-PH. Cdh5-CreERT2;Runx1(flox/flox);SOX17enhKO triple transgenic SRV mice were subjected to mild SuHx treatment. When the SRV mice were treated with corn oil without tamoxifen, they developed significantly elevated RVSP. When the SRV mice were treated with tamoxifen and placed under mild SuHx conditions, they exhibited normal RVSP. **** P < 0.0001.

### Small molecule RUNX1 inhibitors Ro5-3335 and Ro24-7429 reversed SuHx-PH in rats

In the reversal model, rats received RUNX1 inhibition after SuHx-induced PH had already established (Supplementary Figure S3A). SuHx rats treated with vehicle alone developed elevated RVSP compared with Nx control rats (Supplementary Figure S3B). Treatment with the RUNX1 inhibitor Ro5-3335 or Ro24-7429 significantly reduced RVSP in SuHx rats when compared with the vehicle treated SuHx rats, and reversed RVSP to the level comparable to that of the Nx control rats (Supplementary Figure S3B). These results suggest that inhibition of RUNX1 in vivo effectively reverses PH after it has already been established.

### RUNX1 expression is increased in plasma of PAH patients with undetectable levels of SOX17

In order to determine if RUNX1 expression is increased in PAH patients with impaired SOX17 expression, we analyzed gene expression in peripheral blood samples from 359 PAH patients and 72 healthy controls from the UK database. Out of the 359 PAH patients, SOX17 expression was undetectable in 206 and detectable in 153 patients, whereas the expression of RUNX1 could be seen in all subjects. RUNX1 expression was significantly higher in patients with undetectable expression of SOX17 compared to those with detectable SOX17 expression (Figure 11).

**Figure 11.**
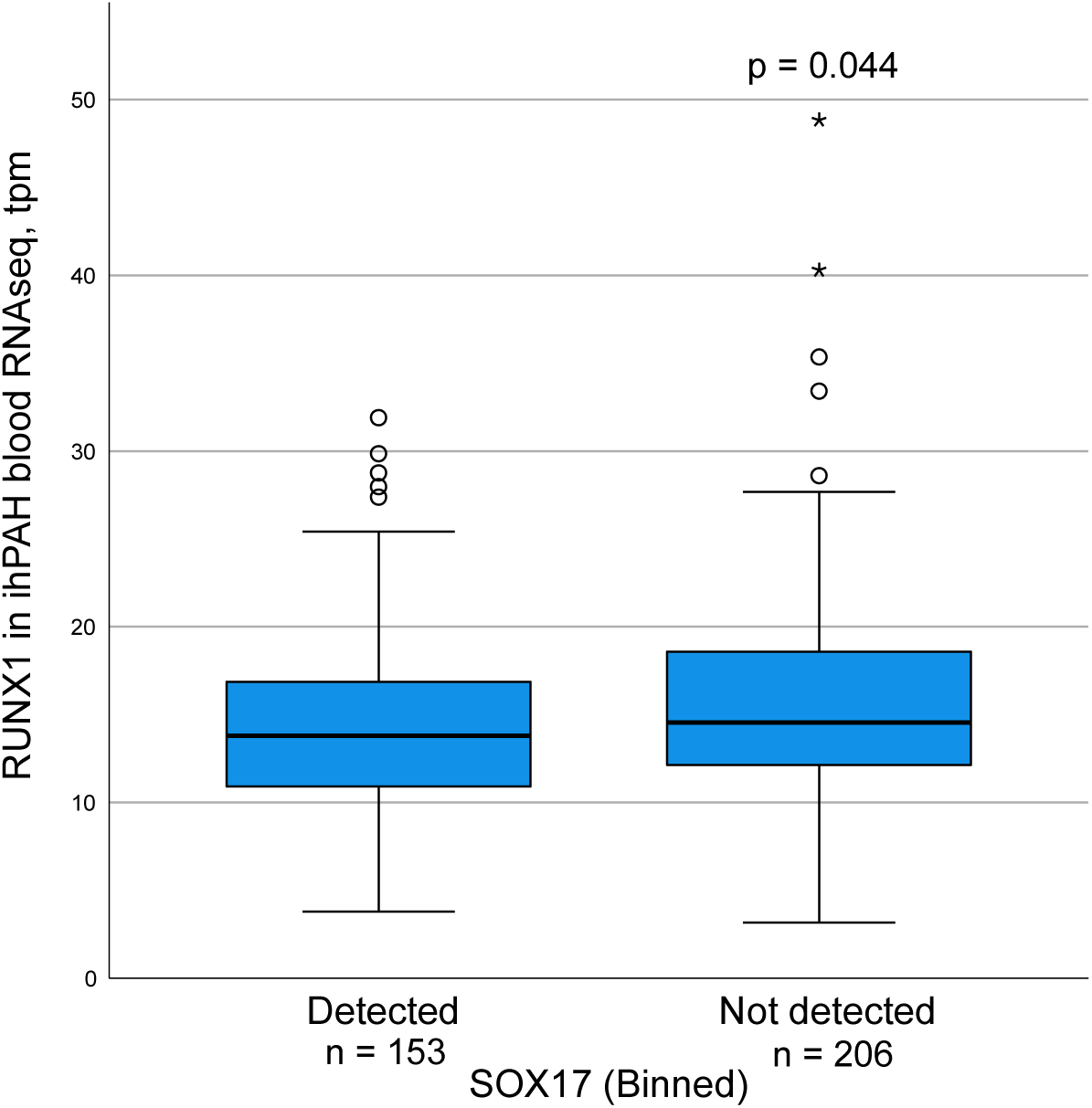
RUNX1 expression is increased in plasma of PAH patients with undetectable levels of SOX17. Out of the 359 PAH patients, SOX17 expression was undetectable in 206 and detectable in 153 patients, whereas the expression of RUNX1 could be seen in all subjects. RUNX1 expression was significantly higher in patients with undetectable expression of SOX17 compared to those with detectable SOX17 expression. * P < 0.05.

## DISCUSSION

Mutations in over 2 dozen genes have been associated with increased risk of PAH and have led to major breakthroughs in our understanding of the pathobiology of PAH. However, evidence for a causal role for many of these genes is lacking. A recent assessment of PAH gene-disease relationships found that SOX17 is one of 12 out of 27 genes with definitive evidence for causal effects for PAH ^31^. SOX17 plays an important role in angiogenesis ^32, 33^ due to its critical role in driving stem and progenitor cells to an endothelial rather than a hematopoietic cell fate ^34^ ^35^. SOX17 directs the expression of numerous genes that are key to maintaining endothelial identity, but its ability to suppress RUNX1 expression by direct binding to specific sites in the RUNX1 promoter has been found to play a crucial role in preventing the differentiation of endothelial cells into hematopoietic stem cells. RUNX1 also promotes the pro-inflammatory macrophage phenotype with increased IL-6 and IL-1β production by enhancing NF-κB activity through direct binding of its subunit p50 ^36^, and favors JAK/STAT signaling through direct transcriptional repression of suppressor of cytokine signaling 3 (SOCS3) ^37,38^. Furthermore, RUNX1 deficiency induces macrophage polarization toward the anti-inflammatory phenotype by enhancing STAT6 phosphorylation ^39^. Importantly, pathogenic roles of both the NF-κB and JAK/STAT signaling pathways have been demonstrated in PAH ^40–42^. Previous work by our group has shown that inhibition of RUNX1 prevents the development of PH and reverses established PH in several animal models of PAH.

In the present study, we sought to determine if impaired SOX17 expression caused endothelial cell dysfunction and pulmonary hypertension by failing to adequately suppress RUNX1. Here we show that SOX17 knockout in HPAEC is associated with marked suppression of numerous EC marker genes and significant impairment of EC function. SOX17 KO HPAECs lost their cobblestone morphologic phenotype and demonstrated decreased apoptosis and significant increases in proliferation and migration and impaired angiogenesis. These changes could also be induced by overexpression of RUNX1 in HPAEC which resulted in downregulation of many of the same EC markers and similar decreases in apoptosis and angiogenesis along with increased proliferation and migration. Notably, SOX17 KO in HPAEC resulted in a marked increase in RUNX1 expression. We then sought to determine if the suppression of EC markers caused by SOX17 KO and RUNX1 overexpression in vitro occurred in EC obtained from PAH patients with SOX17 mutations. Compared to EC derived from iPSCs from healthy controls, EC derived from SOX 17 mutant patient iPSCs had reduced expression of the same EC markers seen in HPAECs with SOX17 KO or RUNX1 overexpression. Importantly, RUNX1 protein levels were significantly increased in EC derived from SOX17 PAH patient IPSCs.

In order to determine if the decrease in EC markers in HPAECs with SOX17 KO and EC derived from SOX17 PAH patient iPSCs is mediated by RUNX1, we treated both sets of cells with the RUNX1 specific inhibitor Ro5-3335. In both HPAECs and iPSC-derived ECs, inhibition of RUNX1 increased the expression of the EC markers that were suppressed. Similarly, knockdown of RUNX1 expression via siRNA also resulted in increased expression of EC markers in the HPAEC SOX17 KO cells and ECs derived from SOX17 mutant PAH iPSCs. Together, these studies strongly suggest that RUNX1 plays an important role in the loss of EC identity that occurs with impaired SOX17 expression in vitro and in patients with PAH associated with SOX17 mutations.

Although SOX17 mutations associated with severe PAH are rare, common SNVs in the SOX17 enhancer region are seen in nearly half of the general population and approximately 3/5 of patients with PAH. Previous studies by our group have shown that common SOX17 SNVs are associated with significant differences in plasma proteins expression and that transgenic mice with deletion of signal 1 of the enhancer region, designed to mimic the common SOX17 SNVs associated with increased risk of PAH in patients, develop more severe hypoxic PH and are more susceptible to SuHx-PH. However, the mechanism by which impairments in the SOX17 enhancer promote PAH is not known and it is unclear if SNVs in the SOX17 enhancer are sufficient to affect RUNX1 expression. Under baseline conditions, we found no difference in RUNX1 expression in whole bone marrow or in the lung between SOX17enhKO and wild-type mice. Interestingly, treatment with low dose sugen and mild hypoxia did not induce PH or increase RUNX1 expression in wild-type mice but did so in SOX17enhKO mice. Treatment of SOX17enhKO mice with 2 different RUNX1 inhibitors completely prevented the increase in RVSP and right ventricular hypertrophy induced by low dose Sugen and mild hypoxia. Prevention of PH in the SuHx-PH susceptible SOX17enhKO mice could also be accomplished by inducible endothelial specific knockout of RUNX1 demonstrating that the inhibitor effect of Ro5-3335 and R24-7245 was due to their effect on RUNX1.

Assessing the expression of transcription factors such as SOX17 and RUNX1 in patients with PAH may be challenging due to the limited supply of tissue samples and the low level of expression in the lung. In earlier studies, we found that plasma RUNX1 expression was increased in PAH patients compared to healthy volunteers. In order to determine if this increase in RUNX1 was related to SOX 17 expression, we analyzed plasma gene expression in the UK PAH database. We found that RUNX1 expression was significantly higher in PAH patients in whom SOX17 expression was not detectable compared to patients with detectable SOX17 expression. The low level of SOX17 expression in plasma samples is not surprising considering that most of the DNA examined is derived from circulating myeloid cells which normally have low levels of SOX17 expression. The low level of SOX17 expression prevented us from correlating SOX17 and RUNX1 expression directly.

Identification of genetic impairments that are associated with increased risk of disease is helpful in furthering the understanding of disease pathology, but advancing the treatment these diseases requires the development of interventions that can mitigate or reverse the cellular mechanisms responsible for pathologic defects caused by altered gene expression. The discovery that mutations in bone morphogenic protein receptor 2 (BMPR2) are responsible for nearly 3/4 of cases of heritable PAH and approximately 1/5 of sporadic PAH cases lead to the development of the recently approved activin receptor 2A (ActRIIA) ligand trap, sotatercept which is designed to rebalance signaling in the TGF-beta pathway affected by impaired BMPR2 expression, This medication has been shown to improve clinical outcomes in PAH patients when added to other PAH medications and appears to be a major advancement in the treatment of PAH (ref). However, clinical trials with sotatercept found multicomponent improvement in 6 MWD, WHO functional class, and NT-proBNP in only 39% of patients compared to 10% of placebo-treated patients ^43^. Furthermore, its use has been associated with a high incidence of adverse off-target effects including erythrocytosis, telengectasias, and pericardial effusions ^43, 44^. These limitations underscore the need for additional therapies to improve efficacy and tolerability.

In conclusion, findings from the present study show that RUNX1 expression is higher in ECs with impaired SOX17 expression and that overexpression of RUNX1 suppresses EC cell markers and induces EC dysfunction. Importantly, both the suppression of EC markers in SOX17 deficient cells and the increased susceptibility to SuHx-PH in SOX17enhKO mice can be reversed by RUNX1 inhibition. These findings identify SOX17/RUNX1 signaling as an important component of PAH pathogenesis and a potential target for novel PAH therapies. Small molecule compound RUNX1 inhibitors such as the ones used in this study have been developed for clinical use and have been found to be highly effective in reversing established PH in several pre-clinical models of PH by our group. Whether or not they may be of particular benefit in patients with rare or common SOX17 mutations remains to be determined. Interestingly, subgroup analysis of clinical trials with sotatercept, found that its clinical benefit was just as effective in patients with idiopathic PAH as in those with heritable PAH, suggesting that therapies designed to compensate for mutations in genes that are important in modulating pulmonary vascular remodeling may be effective treatments for PAH even in those without genetic mutations. The results from this study suggest that targeting SOX17/RUNX1 signaling may be a novel and effect approach for the treatment of PAH.

## ACKNOWLEDGEMENTS

This work was supported in part by the National Institutes of Health NHLBI R01HL158841 (Olin Liang & James Klinger), NHLBI R01HL174007 (Corey Ventetuolo & Olin Liang), NHLBI T32HL160517 (Olin Liang & Bedia Akosman), NIGMS P30GM145500 (Olin Liang), NIGMS P20GM103652 (James Klinger & Olin Liang), by the Brown Physicians Incorporated Academic Assessment Research Awards (Olin Liang & James Klinger), and by the Rhode Island Hospital George and UPMIFA Funds for Hematology and Oncology Research (Olin Liang).

---

**Supplementary Table S1.**

| ID | Cell Type | Sex | Age |
| --- | --- | --- | --- |
| ICZP020 | LCL (Donor #1) | F | 29 |
| ICZP022 | LCL (Donor #10) | F | 35 |
| ICZP029 | LCL (Donor #3) | F | 51 |
| ICZP030 | LCL (Donor #11) | M | 34 |
| ICZP032 | LCL (Donor #13) | F | 69 |

**B**
| ID | PAH Sub-class | Sex | Age at Dx | Ethn. | Nucleotide Change | Amino Acid Change | Type |
| --- | --- | --- | --- | --- | --- | --- | --- |
| 06-116 | APAH-CHD | F | 3 | EUR | c.226A>G | p.(Met76Val) | D-Mis |
| 29-021 | APAH-Portopulm | F | 57 | AMR | c.277C>A | p.(Leu93Met) | D-Mis |
| 24-002 | IPAH | F | 7 | EUR | c.365_366del | p.(Glu122Alafs*39) | FS |
| 28-150 | APAH-CHD | F | 31 | AMR | c.392A>G | p.(Asp131Gly) | D-Mis |
| 05-192 | IPAH | F | 40 | AMR | c.392A>G | p.(Asp131Gly) | D-Mis |
| 06-005 | IPAH | F | 5 | EUR | c.418C>T | p.(Arg140Trp) | D-Mis |
| 06-012 | IPAH | F | 5 | AMR | c.499_520del | p.(Leu167Trpfs*213) | FS |
| 08-008 | DTOX | M | 31 | EUR | c.788del | p.(Pro263Argfs*124) | FS |
| 07-059 | IPAH | F | 66 | EUR | c.1190C>T | p.(Ser397Leu) | D-Mis |

Clinical and genetic features of patients with SOX17 variants included in this study. All variants were annotated using transcript NM\_022454.3. D-Mis indicates damaging missense variants and FS indicates frameshift variants. IPAH, idiopathic pulmonary arterial hypertension; APAH-CHD, associated pulmonary arterial hypertension with congenital heart disease; DTOX, drug/toxin-associated PAH; Ethn., Ethnicity; AMR, Admixed American ancestry; EUR, European ancestry.

**Supplementary Table S2.** List of primary antibodies used in western blots.

| Primary antibody | Isotype | Source | Dilution |
| --- | --- | --- | --- |
| RUNX1 Antibody (A-2) | Mouse | Santa Cruz Biotechnology (sc-365644) | 1:500 |
| CD144 (16B1) | Mouse | Invitrogen (14-1449-82) | 1:250 |
| SOX17 (D1T8M) | Rabbit | Cell Signaling Technology (81778) | 1:1000 |
| PECAM-1 (H-300) | Rabbit | Santa Cruz Biotechnology (sc-8306) | 1:250 |
| Von Willebrand Factor | Rabbit | Abcam (Ab6994) | 1:500 |
| VEGFR2/KDR | Mouse | Proteintech (67407-1-Ig) | 1:1000 |
| CD54/ICAM1 | Rabbit | Cell Signaling Technology (4915S) | 1:1000 |
| CD14 (EPR3653) | Rabbit | Abcam (ab133335) | 1:500 |
| Anti-CD27 [EPR8569] | Rabbit | Abcam (ab131254) | 1:1000 |
| VCAM-1 (SA05-04) | Rabbit | Invitrogen (MA5-31965) | 1:1000 |
| β-ACTIN | Rabbit | Cell Signaling Technology (4967) | 1:5000 |

**Supplementary Figure S1.**
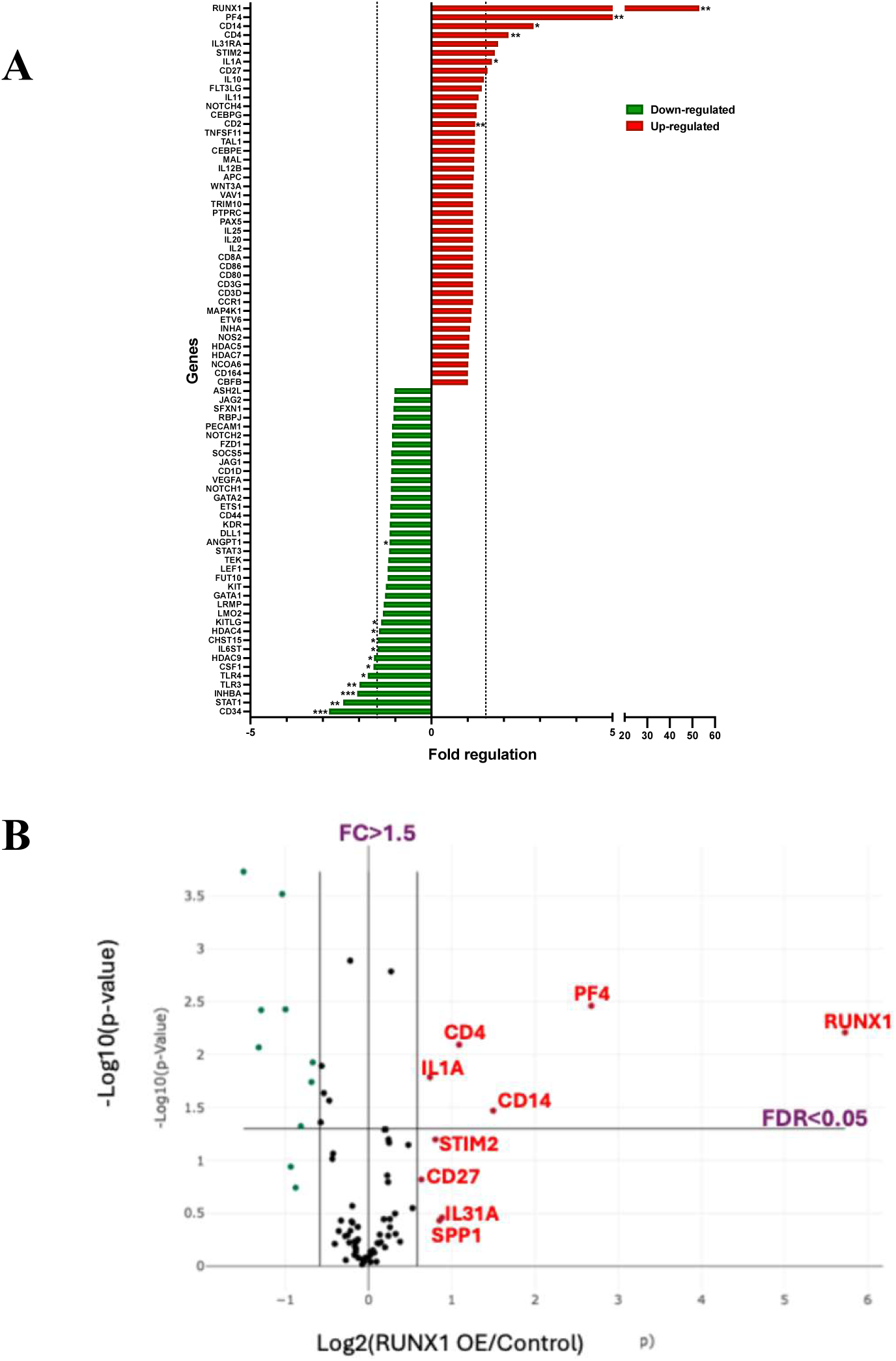
Human Hematopoiesis RT²-PCR Array analyses in HPAECs overexpressing *RUNX1* (*RUNX1 OE*) compared to control vector transduced cells. Gene expression values were normalized to housekeeping genes and represented as fold change relative to controls. Bar graphs (A) and Volcano plots (B) highlight differentially expressed genes meeting the significance criteria of fold change > 1.5 and FDR < 0.05. *p < 0.05, **p < 0.01, ***p < 0.001 indicate statistical significance.

**Supplementary Figure S2.**
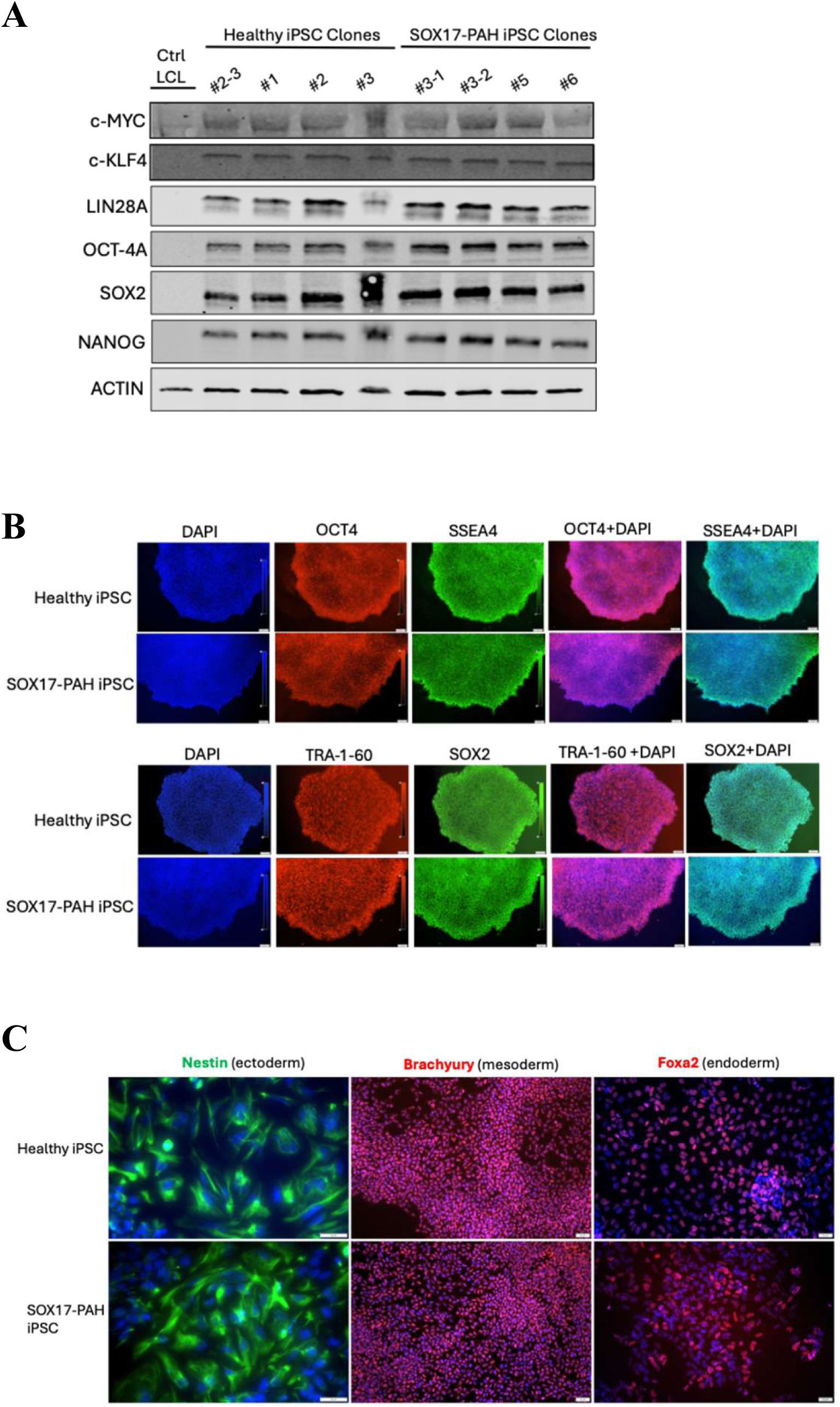
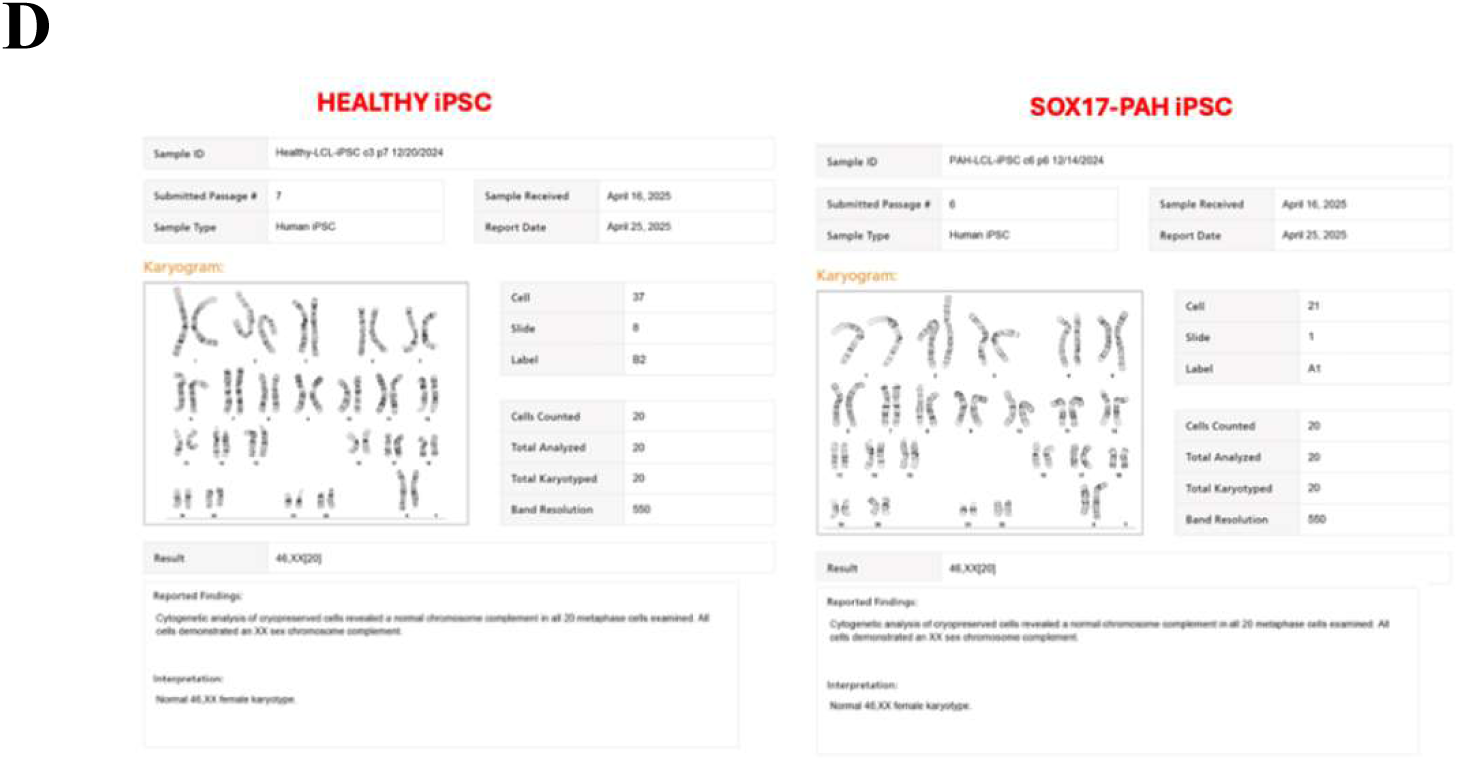
Characterization of iPSCs derived from lymphoblastoid cell lines (LCLs) of healthy controls and SOX17 mutant PAH patients. (A) Whole-cell lysates from control and PAH LCL-derived iPSC colonies were analyzed by western blotting for pluripotency marker expression. Protein levels of c-MYC, KLF4, LIN28A, OCT4A, SOX2, and NANOG were assessed, with ACTIN used as a loading control. (B) Immunocytochemical staining of healthy control and SOX17-PAH LCL-derived iPSC colonies for pluripotency markers OCT4 (red), SSEA4 (green), SOX2 (green), and TRA-1-60 (red). Nuclei were counterstained with DAPI (blue). The rightmost panels show merged images. Scale bar: 50 µm. (C) Trilineage differentiation potential of healthy control and PAH LCL-derived iPSCs assessed by immunocytochemistry. Differentiated cells were stained for ectoderm (Nestin, green), mesoderm (Brachyury, red), and endoderm (FOXA2, red) lineage markers. Scale bar: 20 µm. (D) Representative karyotype analysis of control and SOX17-PAH LCL-derived iPSC clones demonstrating normal chromosomal integrity.

**Supplementary Figure S3.**
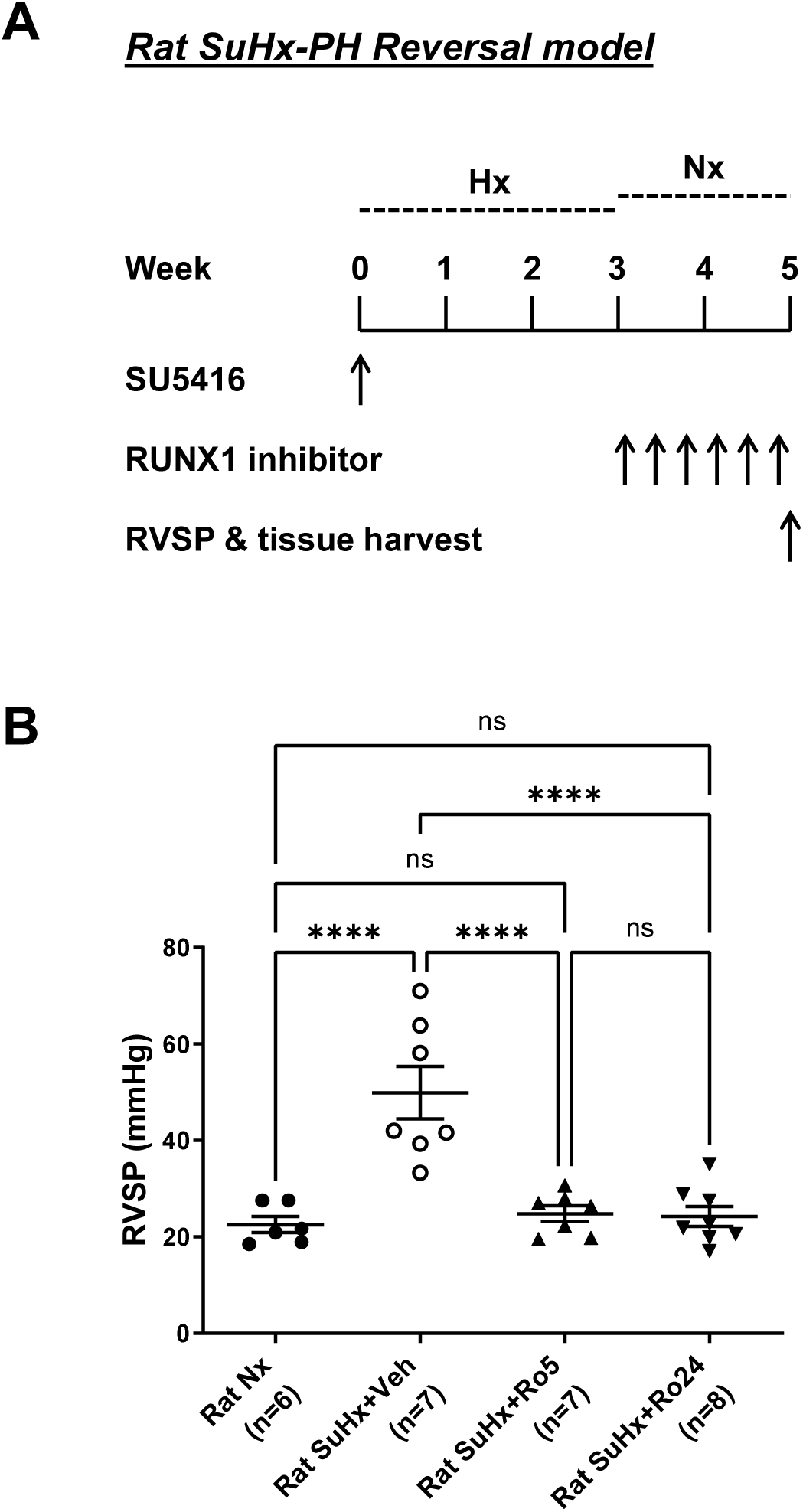
Small molecule RUNX1 inhibitors Ro5-3335 and Ro24-7429 reversed SuHx-PH in rats. (A) Experimental design of a SuHx-PH reversal model in rats. (B) Both Ro5-3335 (20 mg/kg) and Ro24-7429 (40 mg/kg) effectively reversed SuHx-PH in rats. **** p < 0.0001 indicate statistical significance.

